# The influenza A virus endoribonuclease PA-X usurps host mRNA processing machinery to limit host gene expression

**DOI:** 10.1101/442996

**Authors:** Lea Gaucherand, Brittany K. Porter, Summer K. Schmaling, Christopher Harley Rycroft, Yuzo Kevorkian, Craig McCormick, Denys A. Khaperskyy, Marta Maria Gaglia

## Abstract

Many viruses globally shut off host gene expression to inhibit activation of cell-intrinsic antiviral responses. However, host shutoff is not indiscriminate, since viral proteins and host proteins required for viral replication are still synthesized during shutoff. The molecular determinants of target selectivity in host shutoff remain incompletely understood. Here, we report that the influenza A virus shutoff factor PA-X usurps RNA splicing to selectively target host RNAs for destruction. PA-X preferentially degrades spliced mRNAs, both transcriptome-wide and in reporter assays. Moreover, proximity-labeling proteomics revealed that PA-X interacts with cellular proteins involved in RNA splicing. The interaction with splicing contributes to target discrimination and is unique among viral host shutoff nucleases. This novel mechanism sheds light on the specificity of viral control of host gene expression and may provide opportunities for development of new host-targeted antivirals.

## INTRODUCTION

Despite their small genomes, influenza A viruses (IAVs) dedicate multiple proteins to the suppression of host gene expression, or “host shutoff”, which limits host antiviral responses. One of the IAV host shutoff proteins is the endoribonuclease PA-X, which selectively degrades host RNAs (Jagger et al., 2012; Khaperskyy et al., 2016) and limits innate immune responses *in vivo.* PA-X-deficient viruses induce stronger innate immune and inflammatory responses in mice, chickens and pigs (Gao et al., 2015; Gong et al., 2017; Hayashi et al., 2015; Hu et al., 2015, 2016; Jagger et al., 2012; Xu et al., 2017). In some IAV strains, the immune-evasion activity of PA-X reduced inflammation-induced pathology, thereby protecting the host and reducing mortality (Gao et al., 2015; Gong et al., 2017; Hu et al., 2015, 2016; Jagger et al., 2012). While the role of PA-X in immune evasion is well established, its molecular mechanism of action remains poorly understood.

PA-X is produced by ribosomal frameshifting during translation of the IAV PA segment (segment 3) (Firth et al., 2012; Jagger et al., 2012). The frameshift generates a protein that shares the PA amino-terminal ribonuclease (RNase) domain, but has a unique carboxy-terminal sequence known as the X-ORF. The X-ORF is required for PA-X nuclear accumulation and function (Hayashi et al., 2016; Khaperskyy et al., 2016; Oishi et al., 2015). Despite this non-canonical production mechanism, PA-X is encoded by all IAV strains (Shi et al., 2012). We have previously shown that PA-X selectively degrades RNAs transcribed by host RNA polymerase II (Pol II), but not other polymerases (Khaperskyy et al., 2016). Importantly, this characteristic leads to the protection of viral RNAs, because they are generated by the viral RNA-dependent RNA polymerase (RdRp) (Khaperskyy et al., 2016). However, the mechanism for PA-X targeting of Pol II transcripts is not known.

Other viruses also encode host shutoff RNases that selectivity target Pol II transcripts, including vhs from herpes simplex viruses, the SOX/BGLF5 family of proteins in gammaherpesviruses and the severe acute respiratory syndrome-related coronavirus (SARS-CoV) non-structural protein 1 (nsp1) (Covarrubias et al., 2009, 2011; Elgadi et al., 1999; Gaglia et al., 2012; Glaunsinger and Ganem, 2004a; Kamitani et al., 2006; Rowe et al., 2007). However, SARS nsp1 and the herpesviral host shutoff proteins operate in the cytoplasm and only degrade transcripts that are bound by components of the protein synthesis machinery (Covarrubias et al., 2011; Feng et al., 2001; Gaglia et al., 2012; Kamitani et al., 2009). By contrast, PA-X accumulates in the nucleus and translation is neither necessary nor sufficient to direct RNA degradation (Hayashi et al., 2016; Khaperskyy et al., 2016). For example, PA-X cannot degrade a translated reporter mRNA that bypasses canonical nuclear 3’-end processing mechanisms (Khaperskyy et al., 2016). These findings suggest that PA-X has a unique mechanism for targeting host RNAs prior to translation, perhaps in conjunction or concurrently with RNA processing and the assembly of functional messenger ribonucleoprotein (mRNP) complexes. Our previous analysis of select transcripts also suggests that not all Pol II transcripts are equally susceptible to PA-X degradation (Khaperskyy et al., 2016), similar to what has been described for other known viral host shutoff RNases (Glaunsinger and Ganem, 2004b). In agreement with this, a recent study of the host transcriptome during IAV infection showed that certain functional classes of RNAs were spared from shutoff, although no specific link to PA-X activity was established (Bercovich-Kinori et al., 2016).

Here, we profiled transcriptome-wide PA-X targets in human lung A549 cells, in both *de novo* infection and PA-X ectopic expression models. This analysis revealed that PA-X susceptibility was tightly linked to Pol II transcript splicing. PA-X-mediated RNA down-regulation correlated with the number of splice sites on target RNAs, irrespective of RNA length. Moreover, an intronless mRNA reporter was largely PA-X resistant, whereas a matched spliced mRNA reporter was PA-X sensitive. Using proximity labeling proteomics (BioID) (Roux et al., 2012) we identified host proteins involved in mRNA processing that associated with the C-terminal X-ORF, suggesting that PA-X target selection may involve physical interactions with components of the host mRNA processing machinery.

## RESULTS

### PA-X causes global changes in RNA levels during infection

To determine the scope of PA-X specificity for host Pol II transcripts, we profiled RNA levels in cells infected with wild-type (wt) and PA-X-deficient IAV. To generate PA-X deficient mutants in the well-characterized IAV strain A/PuertoRico/8/1934 H1N1 (PR8), we introduced two mutations in the frameshifting site and a nonsense mutation in PA-X, L201Stop, that truncated the X-ORF after 9 amino acids (aa); we dubbed this virus PA(ΔX) (Figure 1A). These mutations were carefully designed to ensure that the PA ORF was not altered. We previously used a strain carrying only the frameshifting mutations, IAV PA(fs) (Figure S1A) (Khaperskyy et al., 2016), but created the IAV PA(ΔX) strain to ensure that any residual frameshifting would produce a non-functional PA-X. We confirmed that the 9-aa truncated PR8 PA-X was largely inactive, as it lost the ability to degrade a β-globin reporter, while a X-ORF truncation to 15-aa had not effect (Fig. 1B). These results are in agreement with previous studies that tested truncations in PA-X variants from other strains (Hayashi et al., 2016; Oishi et al., 2015). We also generated a virus that bore only the L201Stop mutation and dubbed it ‘X9’ (Fig. S1A). We chose to use the PR8 strain because it lacks two other known IAV host shutoff mechanisms, as its NS1 protein does not block host mRNA processing (Das et al., 2008), and its RdRp does not trigger Pol II degradation (Rodriguez et al., 2009).

**Figure 1.**
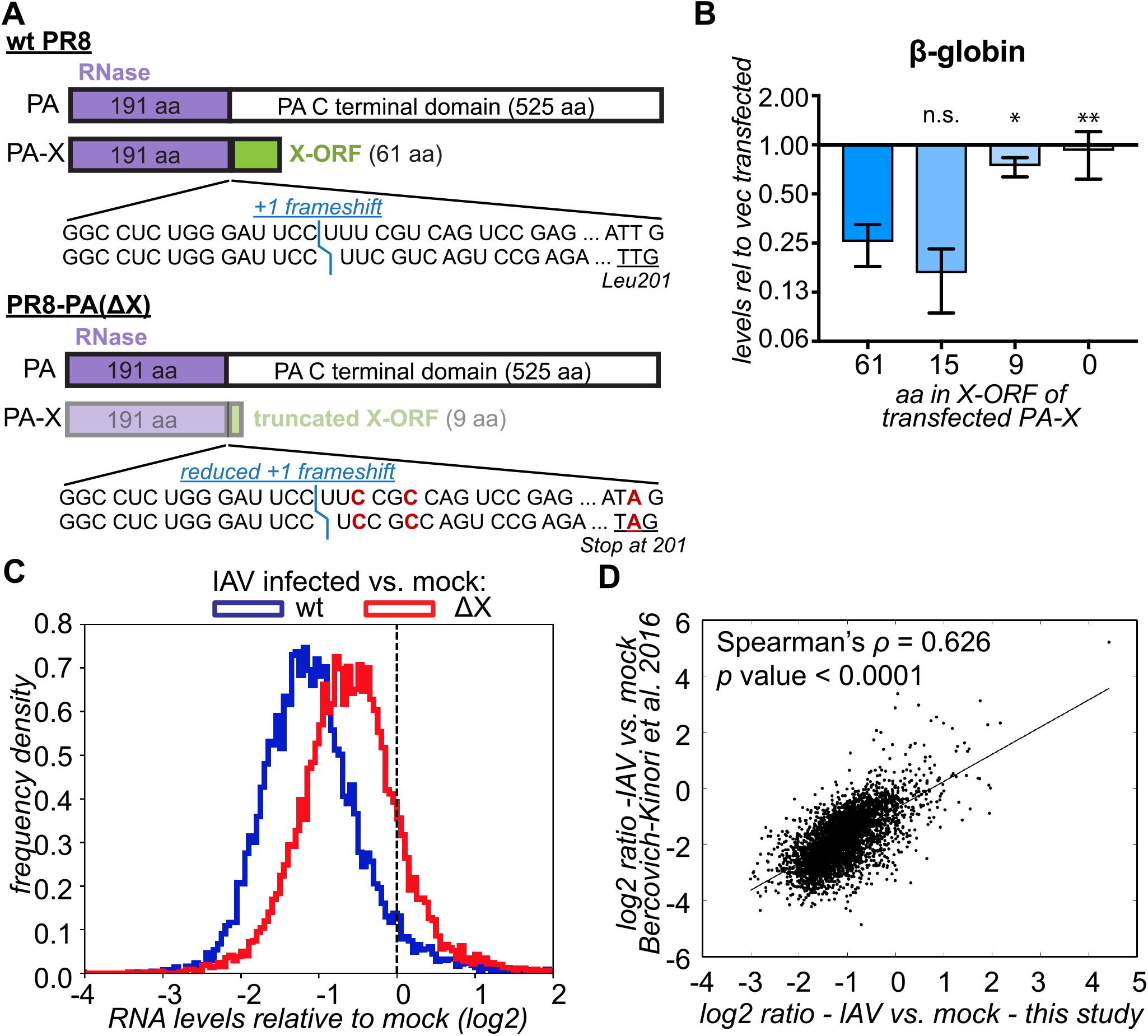
PA-X down-regulates most cellular mRNAs and is a major contributor to host shutoff during influenza A virus infection. (A) Diagram of mutations in the PR8 PA(ΔX) virus. The less intense colors represent lower levels of PA-X. The position of the frameshift is marked in blue. The mutated nucleotides in the frameshifting sequence and at PA-X codon 201 are marked in red. (B) HEK293T cells were transfected for 24 h with a β-globin reporter and wt PR8 PA-X (“61”) or variants with the C-terminal X-ORF truncated after the indicated number of amino acids (aa). The levels of β-globin in PA-X transfected cells were measured by RT-qPCR and are plotted relative to vector transfected cells, after normalization to cellular 18S rRNA. Values represent mean ± standard deviation. N = 3. ns, *, ** = *p* > 0.05, < 0.05, < 0.01, respectively, ANOVA followed by Dunnett’s multiple comparison test *vs.* wt PA-X (61-aa). (C) RNA was collected from infected A549 cells 15 h after infection with wt PR8 or PR8 PA(ΔX) and the levels of all RNAs were measured by RNA-seq. The ratio between the levels in IAV-infected *vs.* mock-infected cells was computed for each RNA and the distribution of the ratios (log2) is plotted as a frequency histogram. The two populations are significantly different (*p* < 0.001) based on the Kolmogorov-Smirnoff test. The dashed line indicated a ratio of 1 (no change). N ≥ 2. (D) The ratio in RNA levels in wt IAV infected cells *vs.* mock-infected cells in our study (15 h post infection, MOI = 1) is plotted against the results from Bercovich-Kinori *et al.* 2016 (8 h post infection, MOI = 5). See also Figures S1, S2.

Using high-throughput RNA sequencing, we found that infection of cells with wt IAV caused a dramatic global decrease in transcript levels compared to mock-infected cells (Figure 1C). However, a small fraction of transcripts escaped shutoff (right tail end of the distribution, Figure 1C). By contrast, shutoff was substantially attenuated in IAV PA(ΔX)-infected cells. Interestingly, the PA(X9) mutant attenuated host shutoff similarly to PA(ΔX) and PA(fs) (Figure S1B). This demonstrates that X-ORF truncation disrupts PA-X function during infection, as predicted from ectopic PA-X expression studies (Hayashi et al., 2016; Khaperskyy et al., 2016; Oishi et al., 2015). Importantly, infection rates by wt and mutant viruses were comparable, based on immunofluorescence staining and viral protein levels (Figures S2A, S2B). We also measured nuclear accumulation of cytoplasmic poly(A) binding protein (PABP), a well-described consequence of host shutoff (Khaperskyy et al., 2014; Kumar and Glaunsinger, 2010; Kumar et al., 2011; Lee and Glaunsinger, 2009). Despite the comparable infection levels, infection with PA(X9) and PA(ΔX) viruses resulted in significantly lower rates of PABP nuclear accumulation compared to wt, confirming impairment of host shutoff (Figures S2A, S2C). Lastly, we compared our RNA-seq results to a previous transcriptome profile of IAV PR8 infected cells (Bercovich-Kinori et al., 2016) and found a strong correlation, despite differences in multiplicity of infection (MOI) and time course of analysis (Figure 1D). Collectively, these data formally demonstrate for the first time that PA-X controls the levels of the majority of RNAs during infection.

### PA-X alone induces global down-regulation of host RNAs

To reduce the complexity of the system, we also examined changes in RNA levels caused by ectopic PA-X expression. We used a doxycycline-inducible PA-X expression system, ‘iPA-X’ cells (Khaperskyy et al., 2016), to induce expression of wt PA-X or the catalytically-inactive D108A mutant. Because the iPA-X cells were clonally selected, we analyzed two independently-generated cell lines for each variant. As expected from previous results with targeted RT-qPCR and metabolic labeling (Hayashi et al., 2015; Jagger et al., 2012; Khaperskyy et al., 2016), wt PA-X robustly down-regulated steady-state transcript levels (Figure 2A). The degree of host shutoff was correlated with the levels of PA-X in the cells (percentage of total reads mapping to PA-X: wt #1= 0.005%-0.006%, wt #10= 0.023%-0.026%), and was dependent on RNase activity, because expression of the PA-X catalytic mutant had no effect (Figure 2A). Importantly, a substantial minority of transcripts were unaffected by PA-X expression (Figure 2A, right tail end of the distributions). Furthermore, we observed correlation between the PA-X-dependent down-regulation of RNAs in the ectopic PA-X expression system and in virus-infected cells (Figure 2B). This indicates that PA-X largely targets the same RNAs in the absence of other viral proteins, and that the ectopic expression model captures the contribution of PA-X to host shutoff during infection. To further validate these findings, we selected representative RNAs from the RNAseq dataset and measured their levels by RT-qPCR. We chose mRNAs that were strongly down-regulated (GAPDH, G6PD) or largely unaffected (HNRNPA0, TAF7, EPC1) both in infected cells and in the ectopic overexpression model. We found that the RT-qPCR results agreed with the RNAseq data in terms of the selective effects on the tested transcripts (Figure 2C). Moreover, the mRNA levels were similarly affected by overexpression of PA-X from the A/Udorn/72 H3N2 (Udorn) strain, suggesting that target selection by PA-X is conserved between different virus strains (Figure 2D). Collectively, these data demonstrate that PA-X broadly targets RNA for degradation, while a subset of RNAs remains unaffected.

**Figure 2.**
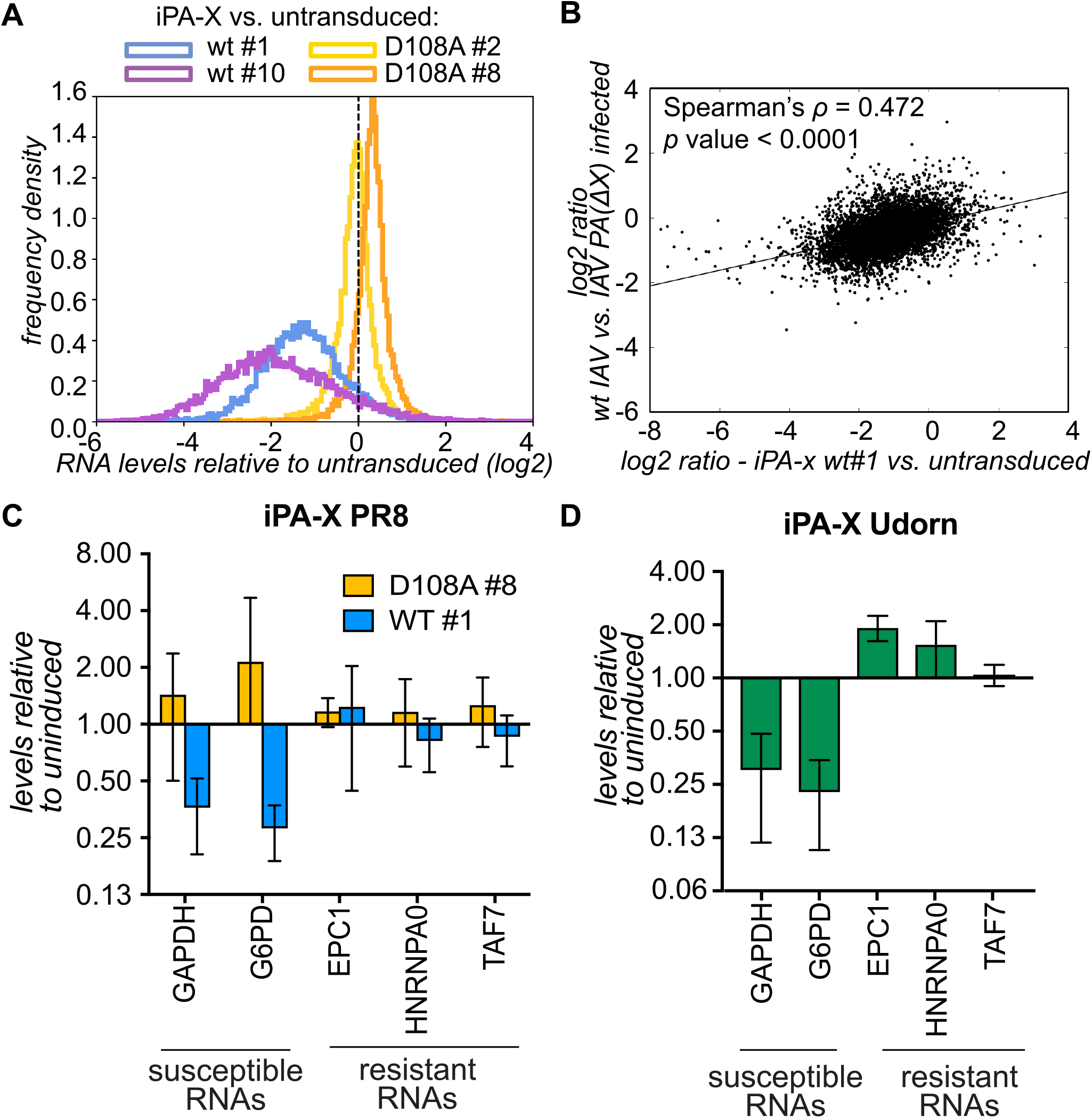
PA-X down-regulates most cellular mRNAs in the absence of other viral proteins. RNA was collected from control (untransduced) A549 cells, or A549 cells expressing doxycycline-inducible PR8 PA-X (wt), PR8 PA-X catalytic mutant (D108A) or Udorn PA-X 18 h after addition of doxycycline. (A) The levels of all RNAs were measured by RNA-seq. The ratio between the levels in PA-X-expressing *vs.* control cells was computed for each RNA and the distribution of the ratios is plotted as a frequency histogram. Two clonal lines were tested for each PA-X variant. The dashed line indicates a ratio of 1 (no change). N ≥ 2. (B) The PA-X-dependent changes in RNA levels in infected cells (ratio in PR8-PA(ΔX) *vs.* wt PR8) are plotted against the changes in cells expressing PR8 PA-X *vs.* control cells. (C-D) The levels of several endogenous mRNAs were measured by RT-qPCR. After normalization to 18S, mRNA levels are plotted relative to uninduced cells. N ≥ 3.

### Specific functional classes of host RNAs are differentially sensitive to PA-X

Although most RNAs were down-regulated by PA-X, the levels of ~25% of RNAs remained largely unchanged (Figures 1C, 2A). To identify these differentially-regulated RNAs, we used *k*-means clustering to group RNAs with similar patterns of regulation (Gasch and Eisen, 2002). Clustering was carried out based on the relative RNA levels in PA-X-overexpressing *vs.* control cells or IAV-infected *vs.* mock-infected cells in the eight datasets we collected (Figures 1C, S1B, 2A). Two sets of RNAs, comprising 55% of the RNAs that were detected in all conditions, were identified as true PA-X targets (Figure 3A). They were down-regulated in a PA-X-dependent manner both during infection and by PA-X ectopic expression. The first group was completely PA-X-specific, because their levels were largely unchanged in IAV PA(ΔX) infected cells (Figure 3A, left panel). The RNAs in the second group were PA-X-sensitive, but were partially down-regulated by other mechanisms during IAV PA(ΔX) infection (Figure 3A, right panel). By contrast, 28% of the RNAs were host shutoff resistant and were not down-regulated by infection or PA-X expression (Figure 3B). In addition, the *k-*means algorithm identified a group of RNAs that were down-regulated during infection by a PA-X-independent mechanism, and yet were PA-X-sensitive in PA-X-expressing cells (Figure 3C). The levels of these transcripts may be substantially decreased by other regulatory mechanisms during infection, such that their targeting by PA-X is masked. Interestingly, based on gene ontology (GO) term analysis, the host shutoff-resistant RNAs were significantly enriched for genes involved in transcription and translation, including ribosomal RNA processing, ribosomal proteins, and membrane protein synthesis (Figure 3D). This result is consistent with the IAV requirement for host biosynthetic machinery and previous observations by Bercovich-Kinori *et al.* (Bercovich-Kinori et al., 2016). Collectively, these results suggest that while PA-X can target many RNAs, it retains some specificity for functional classes of RNAs.

**Figure 3.**
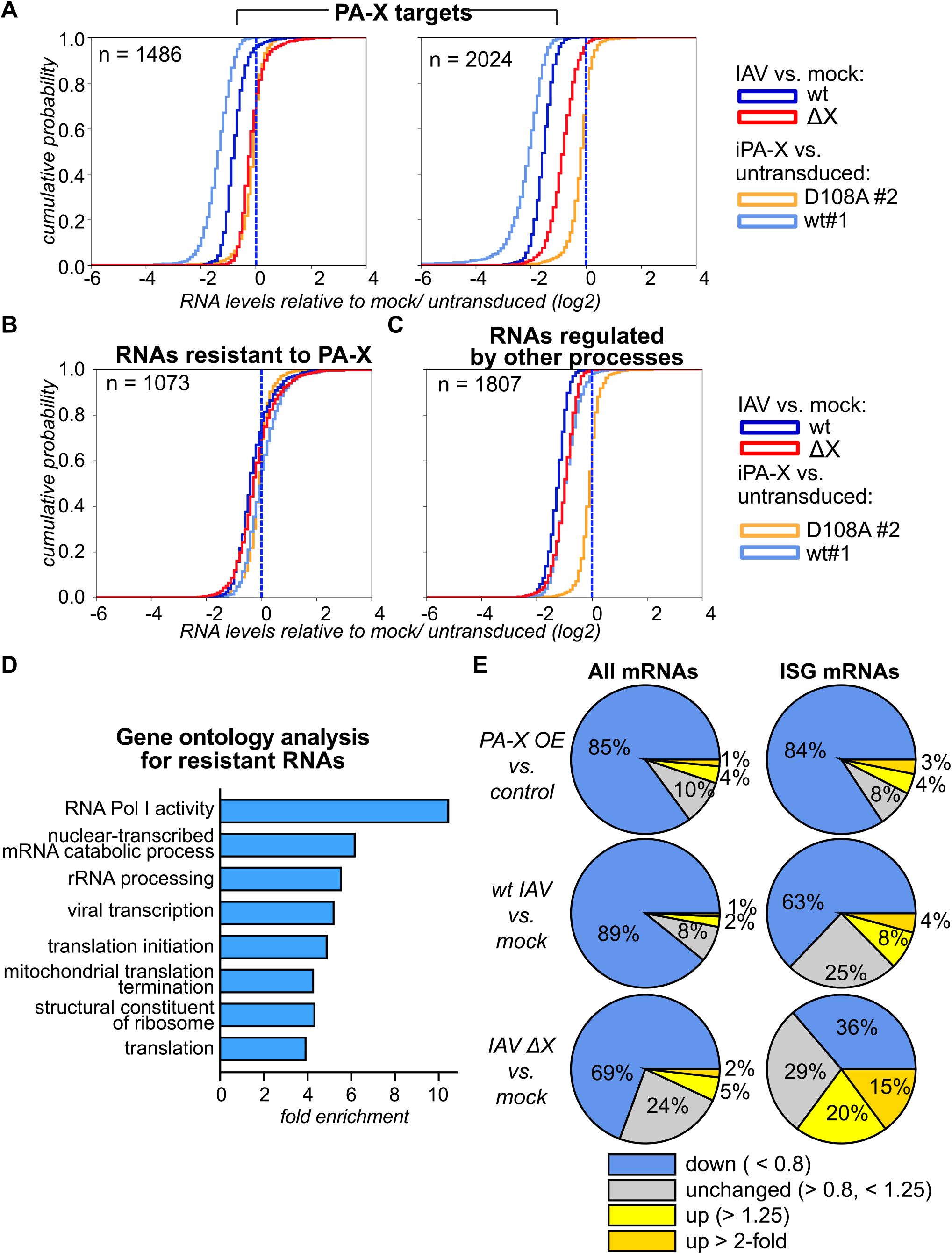
*k*-means clustering reveals differentially regulated groups of mRNAs. (A-C) Cluster3 was used to divide the cellular mRNAs in four clusters based on the pattern of fold changes in iPA-X cells (PA-X wt or D108A catalytic mutant *vs.* control) and in cells infected with IAV (wt or PA-X-deficient IAV *vs.* mock). Cumulative probability histograms of the fold changes for each of the classes are plotted: A: two groups of PA-X targets; B: RNAs that are resistant to PA-X; C: potential PA-X targets that are regulated by other processes during infection. All the datasets collected were used for clustering, but only select datasets are plotted for simplicity. (D) DAVID was used to identify overrepresented gene ontology (GO) terms for biological processes and molecular functions among RNAs that are resistant to PA-X. The fold enrichment is plotted for GO terms that had corrected *p* values < 0.01. (E) Pie-charts showing the percentage of genes that are up-and down-regulated in infected cells and cells expressing PA-X. Left: all RNAs detected in the RNAseq (wt *vs.* mock: N = 8573, PR8-PA(∆X) *vs.* mock: N = 8848, PA-X overexpressing (OE) *vs.* control: N = 8554); Right: interferon-stimulated genes only (ISGs; wt *vs.* mock: N = 167, PR8-PA(∆X) *vs.* mock: N = 168, PA-X overexpressing (OE) *vs.* control: N = 159).

In addition to this unbiased analysis, we examined how PA-X expression affected the levels of interferon-stimulated genes (ISGs), which are induced in infected cells and function in intrinsic antiviral defense (Schoggins et al., 2011). We observed that although ISGs were induced during IAV infection, as demonstrated by their higher expression compared to all detected RNAs, their levels were even higher in the absence of PA-X (Figure 3E). While only 12% of ISGs were up-regulated during wt IAV infection, and only 4% were induced more than two-fold, 35% of the ISGs were up-regulated in IAV PA(ΔX)-infected cells, with 15% above two-fold. While the activity of PA-X is clearly not limited to ISGs, these data indicate that PA-X contributes to dampening the cell-intrinsic response to infection.

### PA-X strongly and preferentially down-regulates spliced Pol II transcripts

We have previously shown that PA-X preferentially degrades RNA transcribed by Pol II (Khaperskyy et al., 2016). A key difference between transcripts of Pol II and other cellular Pols is that Pol II RNAs can be spliced. The process of splicing is mechanistically linked to transcription, because recruitment of the spliceosome is mediated by the C-terminal domain of the large subunit of Pol II (Gu et al., 2013). However, a subset of Pol II transcripts naturally lack introns. When we separately analyzed spliced *vs*. intronless RNAs, we found that PA-X down-regulated spliced RNAs more than intronless RNAs (Figure 4A). Our targeted validation also showed that two intronless mRNAs, TAF7 and HNRNPA0, were not down-regulated by PA-X (Figure 2C). Moreover, during infection, down-regulation of spliced RNAs was clearly dependent on PA-X, whereas most of the down-regulation of intronless RNAs was PA-X-independent (Figure 4B). For these analyses, we only included intronless RNAs longer than 300 nt. This procedure enriched for mRNAs and long non-coding RNAs, excluding small non-coding RNAs like microRNAs and small nuclear and nucleolar RNAs, and ensured that the length distribution was similar between spliced and intronless RNAs. We also analyzed how the number of exons affected degradation, because the number of splice sites varies dramatically among spliced RNAs. Interestingly, there was a significant negative correlation between the number of exons in a transcript and its steady-state levels in PA-X-expressing and infected cells (PA-X-expressing cells: Spearman’s ρ = −0.52, Figure 4C; IAV-infected cells: ρ = −0.47, Figure S3A). This result suggests that RNAs with more exons are more susceptible to PA-X degradation. However, the number of exons in an RNA is often proportional to RNA length and a prior study reported a relationship between IAV host shutoff and transcript length (Bercovich-Kinori et al., 2016). We also observed a correlation between degradation and RNA length (ρ = −0.38 for both PA-X-expressing, Figure 4D, and IAV-infected cells, Figure S3B). To determine whether the exon number or the transcript length was important, we examined RNAs of similar length or with a specific number of exons. We still found a robust negative correlation between RNA degradation and exon number among RNAs of similar length (length = 3.5-4.0 kb, ρ = −0.42, N = 674, Figure 4E). Similar correlations were also seen for subsets of RNAs of other lengths (Figure S3C). By contrast, there was only a small correlation between degradation and RNA length among RNAs with the same number of exons (number of exons = 6, ρ = −0.16, N = 642, Figure 4F). Again, similar correlations were seen for other exon numbers (Figure S3D). We also found that there was no correlation between the degradation and the GC content of RNAs in our dataset (Figures S3E, S3F).

**Figure 4.**
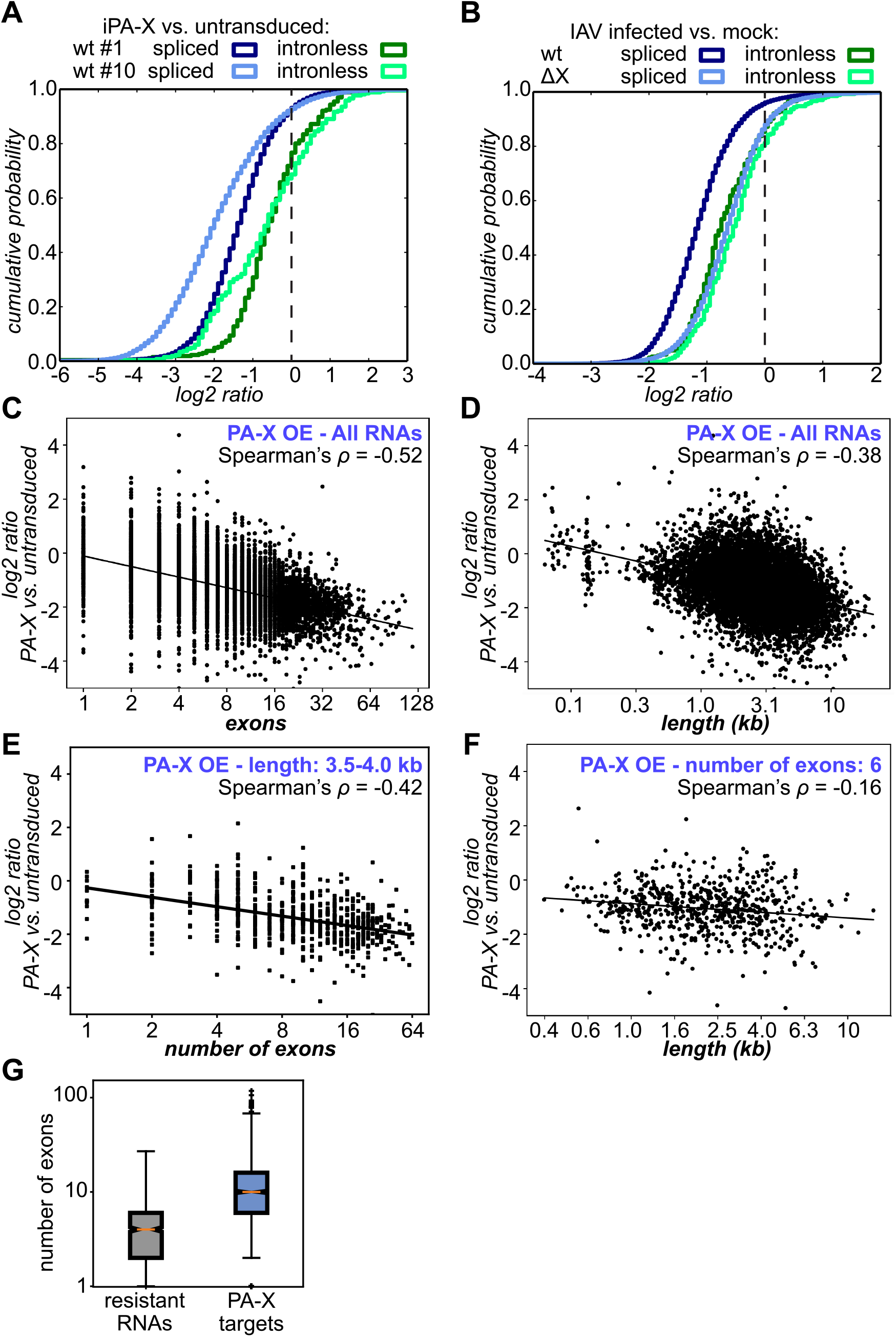
RNAs that are not spliced are less sensitive to regulation by PA-X. (A-B) The RNAseq results from Figures 1C, 2A are plotted separately for spliced and intronless RNA as a cumulative distribution histogram. A: cells overexpressing wt PA-X, clones #1 and #10; B: cells infected with wt PR8 *vs.* PR8-PA(∆X). (C-F) The relative RNA levels in PA-X overexpressing (“PA-X OE”, clone #1) *vs.* control cells are plotted against the number of exons (C, E, log2 scale) or transcript length in kb (D, F, log10 scale). For C and D all RNAs were used, whereas for E and F only RNAs that had 6 exons (E) or were 3.5-4.0 kb long (F) were used. All correlation based on Spearman’s test are statistically significant (*p* < 0.001). (G) The number of exons for RNAs identified in the clustering analysis in Figure 3 is plotted. The two groups of PA-X targets from Figure 3A are plotted together. *p* < 0.01, Kolmogorov-Smirnoff test. See also Figure S3.

The results from the clustering and GO analysis (Figures 3A-D) suggested that PA-X differentially regulates RNAs from specific functional groups, while the analysis in Figures 4A-F suggested a difference based on the structure of the nascent transcript. Interestingly, there was a connection between the structural and functional specificity. RNAs classified as resistant by *k-*means clustering had fewer exons than those that were classified as PA-X targets (Figure 4G). Collectively, these results suggest that targeting by PA-X is connected to mRNA splicing, and that the preference based on splicing has consequences for the selection of functionally relevant targets by PA-X.

### Splice sites confer susceptibility to PA-X

Endogenous RNAs with different numbers of exons also have different sequences, lengths and post-transcriptional modifications. To investigate the effect of splicing in a more controlled system, we examined the same RNA in both spliced and intronless forms. We cloned the cDNA and the full genomic sequence for interferon λ2 (IFN-λ2, Figure 5A), a type III IFN that contributes to IAV immune responses (Jewell et al., 2010). In our RNAseq dataset, IFN-λ2 transcripts were only detectable in IAV-infected cells, and were higher in cells infected with the IAV PA(ΔX) virus (Figure S4A), suggesting that PA-X down-regulates IFN-λ2 mRNA in infected cells. We co-transfected the IFN-λ2 expression vectors with PR8 PA-X and a luciferase construct containing an intron (Younis et al., 2010), which served as a positive control for PA-X activity. We confirmed that IFN-λ2 mRNA was expressed and exported into the cytoplasm at similar levels irrespective of the construct used (Figures S4B, S4C). We also checked that the five introns were properly spliced by PCR analysis across the splice sites (Figures 5E, S4D). When we analyzed the effect of PA-X on the two mRNAs, we found that PA-X down-regulated the IFN-λ2 mRNA expressed from the full genomic region (Figure 5B). However, the levels of the same IFN-λ2 mRNA expressed from an intronless cDNA construct were only minimally reduced (Figure 5B). In both conditions, the control luciferase reporter was down-regulated by PA-X. PA-X proteins from the 2009 pandemic H1N1 strains (such as A/California/7/09(H1N1) (CA/7) and A/Tennessee/1-560/2009) and the Udorn H3N2 strain also preferentially degraded intron-containing target RNAs (Figure 5C). These results confirm the prediction based on our RNAseq data that splicing is important for PA-X substrate recognition. In addition, we tested the down-regulation of IFN-λ2 mRNAs expressed from chimeric constructs that contained only one of the five introns from the original genomic sequence, to determine whether one splicing event was sufficient to restore PA-X targeting. Indeed, addition of single introns increased susceptibility to PA-X (Figure 5D). Interestingly, not all introns were effective, in part because of variable splicing efficiency. For example, intron 4 was not spliced efficiently in the absence of other introns and also did not restore PA-X susceptibility (Figure 5E). This result further strengthens the link between splicing and PA-X susceptibility.

**Figure 5.**
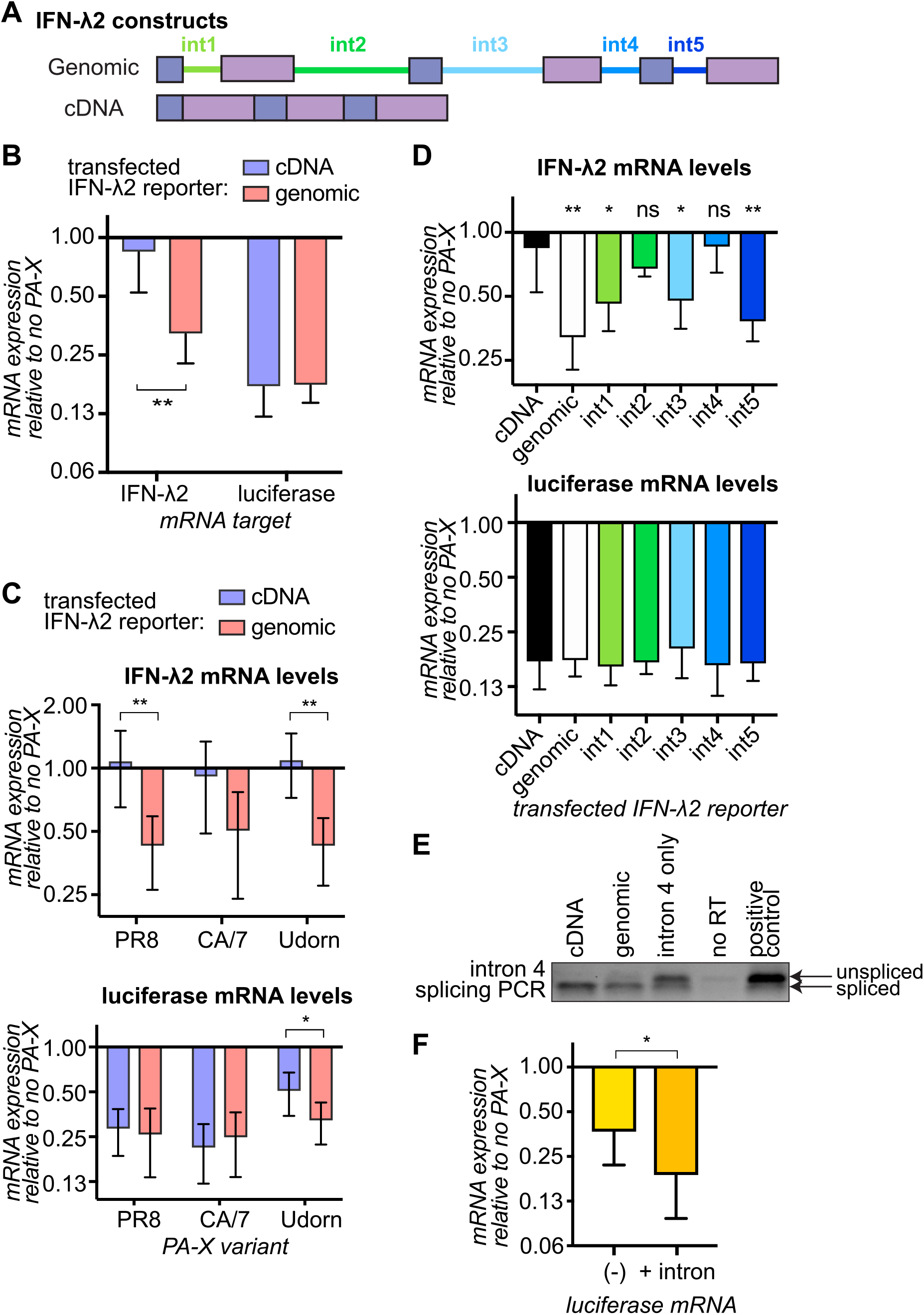
Addition of introns and splicing events promotes degradation by PA-X. (A) Diagram of IFN-λ2 constructs. Int = intron. (B-E) HEK293T cells were transfected for 24 hrs with reporters expressing a luciferase control mRNA and IFN-λ2 mRNA from cDNA, the genomic locus or cDNA with one of the five IFN-λ2 introns added back (D only). Cells were also transfected with PA-X (PR8 variant in B and D, PR8, CA/7 and Udorn variants in C) or vector. The levels of luciferase and IFN-λ2 mRNAs were measured by RT-qPCR and plotted as relative levels in PA-X expressing cells *vs.* vector-transfected cells, after normalization to 18S rRNA. In E, the cDNA from control cells was PCR amplified across intron 4 to test splicing of intron 4. The amplified PCR products are shown (gel image is representative of four experiments). A 1:1 mix of the constructs for IFN-λ2 cDNA and genomic served as a control, to check that both products could be simultaneously amplified. (F) Cells were transfected with an intronless (“(-)”) or an intron-containing (“+ intron”) luciferase reporter and PR8 PA-X. Luciferase RNA levels were measured by RT-qPCR, normalized by 18S rRNA and are plotted relative to vector transfected cells. Values in this figure represent mean ± standard deviation. *,**: *p* < 0.05, 0.01, B, C: ANOVA followed by Tukey’s pairwise test; D: ANOVA followed by Dunnett’s test, *p* values relative to cDNA construct-transfected cells; F: Student’s *t* test. N ≥ 4, all panels. See also Figure S4.

Despite these results linking mRNA splicing and PA-X degradation, one unresolved issue is that we and others have previously used intronless reporters to study PA-X, and these reporters appeared to be efficiently degraded. To resolve this issue, we compared the fate of matching intronless and spliced luciferase reporters (Younis et al., 2010). The spliced reporter, which we used as a control in Figures 5B-D, contains a portion of the β-globin intron (Younis et al., 2010). Interestingly, PA-X had a more robust effect on the spliced mRNA, although it could also down-regulate intronless luciferase mRNA (Figure 5F). These results suggest that addition of a single splicing event further promotes degradation by PA-X. Reporters are selected for their robust expression, although they are somewhat different from endogenous cellular mRNAs. Therefore, it is possible that even intronless reporters associate with cellular factors that are important for PA-X targeting. Nevertheless, our findings recommend the use of intron-containing reporters for future cell-based studies of PA-X.

### The X-ORF mediates interaction with proteins involved in RNA metabolism

As shown in previous studies (Hayashi et al., 2016; Khaperskyy et al., 2016; Oishi et al., 2015) and the RNAseq results with the PR8-PA(X9) virus (Figure S1B), the C-terminal X-ORF is required for PA-X activity. We hypothesized that the X-ORF interacts with cellular proteins that mediate the association of PA-X with target mRNAs, especially in light of our results connecting PA-X targeting with splicing (Figs. 4-5) and mRNA 3’-end processing (Khaperskyy et al., 2016). To identify cellular X-ORF-interacting proteins we used a proteomic technique called BioID, which relies on non-specific proximity biotinylation of lysine residues by a modified *E. coli* biotin ligase, BirA* (Roux et al., 2012). Because there are two major classes of PA-X isoforms that differ in X-ORF length (Shi et al., 2012), we fused BirA* to X-ORFs representative of each class: the 61-aa PR8 X-ORF (X61) and the 41-aa CA/7 X-ORF (X41) (Figure 6A). As negative controls, we used BirA* alone and BirA* fused to a mutated version of the PR8 X-ORF in which four positively charged residues were substituted by alanine (X61(4A)). These mutations prevent nuclear localization of a GFP-X-ORF fusion and disrupt mRNA degradation by full length PA-X (Khaperskyy et al., 2016). As expected from our previous studies with GFP fusion and epitope-tagged proteins (Khaperskyy et al., 2016), fusion to the wt X-ORFs, but not X61(4A), led to accumulation of BirA* in the nucleus (Figure S5A). Moreover, BirA* alone and BirA*-X-ORF fusion proteins efficiently biotinylated numerous cellular proteins (Figure S5B).

**Figure 6.**
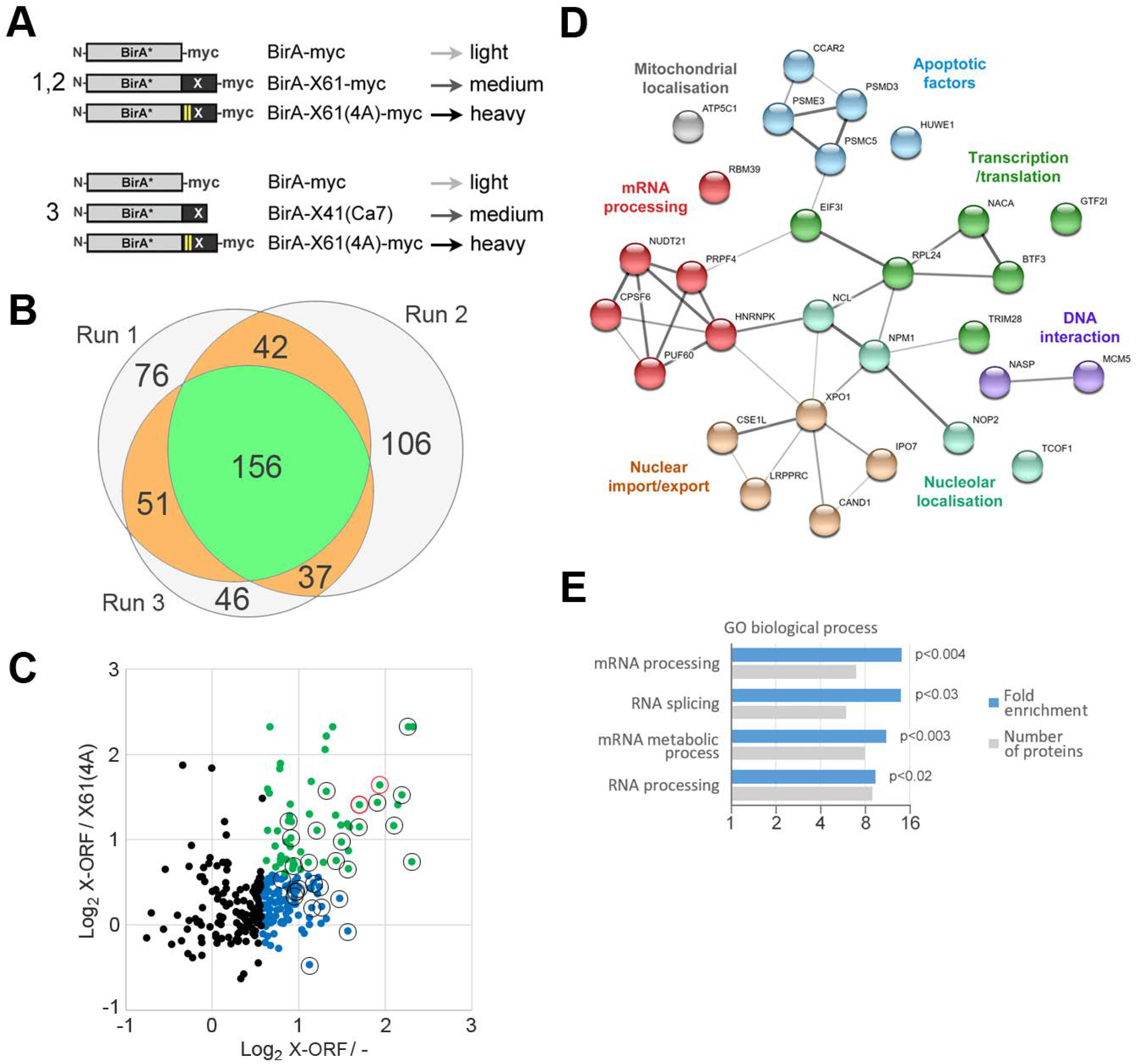
The X-ORF interactome is enriched for proteins involved in mRNA processing. (A) Schematic diagram of BirA* fusion baits used in BioID analysis of X-ORF interacting proteins. Numbers indicate independent runs employing each construct set, with BirA*-myc and BirA*-X61(4A)-myc as negative controls. Light, medium, and heavy refer to light, medium, or heavy isotope tags. (B) Overlap between proteins identified by mass spectrometry by at least two unique peptides in the three BioID runs. (C) Average relative abundance of 286 proteins identified in at least two BioID experiments plotted as log2 ratio of medium *vs.* light (x axis, X-ORF / −) and medium *vs.* heavy (y axis, X-ORF / X61(4A). Green dots represent proteins with >1.5-fold enrichment over both negative controls, blue dots represent proteins with >1.5-fold enrichment over BirA*-myc alone. Black and red open circles indicate proteins that were designated as high-confidence hits (>2.0-fold enriched over BirA*-myc in at least 2 experiments or >1.5-fold enriched over BirA*-myc and BirA*-X61(4A)-myc in all three experiments). Red open circles highlight the two proteins, nucleolin (NCL) and nucleophosmin (NPM1), that were consistently enriched >2.0-fold over both negative controls in all three experiments. (D) STRING protein-protein interaction network for X-ORF BioID high-confidence hits. Apparent nodes were differentially colored (annotations may overlap for individual proteins, only one grouping for each protein was shown for simplicity). (E) Gene ontology (GO) enrichment analysis of X-ORF BioID hits identified by mass spectrometry (black and red circles in C). Fold enrichment and number of proteins are plotted for GO terms with *p* < 0.05. See also Figure S5.

To identify cellular proteins that bind both the X61 and X41 X-ORFs, we collected protein lysates from HEK293T cells expressing the BirA*-X-ORF fusion proteins. We prepared affinity-purified biotinylated proteins for quantitative mass spectrometry, using reductive dimethylation to introduce stable epitope tags. We performed three independent replicates of the experiment, two with BirA*-X61 and one with BirA*-X41, comparing them in each case to the BirA* alone and BirA*-X61(4A) controls (Figure 6A). We identified 156 candidate X-ORF interacting proteins that were represented by at least two unique peptides (Table S5), and were identified in all runs (Figure 6B). Among the 156 candidates, we identified 29 high-confidence interacting proteins based on relative peptide abundance in the test conditions (BirA*-X61 or X41) *vs.* the control conditions (BirA*-X61(4A) or BirA* alone) (Figure 6D, Table S5). These high-confidence hits were enriched more than two-fold over controls in at least two experiments or more than 1.5-fold in all three experiments. Figure 6C depicts the relative peptide abundance of proteins in X61 samples compared to the X61(4A) control or BirA* alone control, with black and red open circles identifying the high confidence hits. The red circles indicate two proteins (nucleolin, nucleophosmin) that were enriched >2-fold compared to controls across all three runs. Because nucleolin and nucleophosmin are abundant proteins that move in and out of nucleoli, these interactions may explain the apparent nucleolar accumulation of biotinylated proteins (Figures S5A, S5C). STRING protein-protein interaction network analysis of the hits revealed several physical and functional interaction nodes, including protein trafficking, transcription/translation and mRNA processing (Figure 6D). Similarly, GO term analysis revealed a strong association with mRNA processing, RNA splicing and mRNA metabolic process functions among the high-confidence hits (Figure 6E). In particular, several proteins associated with mRNA splicing (RBM39, PUF60, PRPF4) and/or polyadenylation (NUDT21/ CPSF5, CPSF6) were identified as X-ORF-interacting proteins. These data show that the PA-X C-terminal X-ORF is physically recruited to protein complexes involved in nuclear mRNA processing, which likely explains its preferential degradation of RNAs that have undergone co/post-transcriptional processing.

## DISCUSSION

A thorough understanding of the molecular mechanism of action of PA-X is required to determine how it selectively degrades host RNAs and contributes to the control of the innate immune response. In this study, we discovered a key aspect of the PA-X mechanism of action: its selectivity for transcripts that have undergone splicing. Our transcriptomic results show that, as expected, PA-X down-regulates many host RNAs, both on its own and in the context of infection. However, some RNAs are less susceptible or completely resistant to PA-X activity. This difference is due in part to the fact that they are either intronless or have fewer introns. Moreover, based on proteomic analysis, the X-ORF of PA-X physically interacts with many proteins involved in cellular RNA metabolism. These data point to a model whereby PA-X associates with RNA metabolism proteins, and this association brings PA-X to target RNAs, perhaps during transcription or early processing. Those RNAs that are not canonically processed, including viral RNAs, are thus not targeted.

Our results confirm that the PA-X-dependent down-regulation of host protein production (Hayashi et al., 2015; Jagger et al., 2012) is due to a PA-X-driven reduction in RNA levels. Moreover, our transcriptomic studies with PA-X-deficient viruses defined both a PA-X-dependent and a PA-X-independent component of RNA down-regulation during IAV infection. The PA-X-independent component is likely due to a recently described generalized reduced transcription of host genes (Bauer et al., 2018), because other modalities of IAV host shutoff are not active in the PR8 strain (Rodriguez et al., 2009; Salvatore et al., 2002). The PR8 NS1 protein does not block 3’-end RNA processing (Das et al., 2008), and its RdRp does not trigger degradation of Pol II (Rodriguez et al., 2009). The mechanism of reduced host transcription in IAV infected cells is not yet known, but it is not dependent on PA-X, because it also occurs during infection with influenza B viruses (Bauer et al., 2018), which do not encode PA-X (Shi et al., 2012). Lastly, clustering analysis revealed that some functional classes of RNAs are spared from PA-X degradation, which agrees with previous results from Bercovich-Kinori *et al.* (Bercovich-Kinori et al., 2016). In particular, mRNAs for proteins involved in translation appear to be spared (Figure 3D, (Bercovich-Kinori et al., 2016)). Our new results suggest a potential explanation for this phenomenon, which is linked to the small number of exons of these mRNAs, particularly RNAs for ribosomal proteins.

The key unexpected finding from our study is the link between PA-X and splicing. Viral host shutoff RNases mostly act at some stage of mRNP loading into the translation apparatus. For example, alpha-herpesvirus vhs directly interacts with translation initiation factors, and it is generally believed that these interactions mediate RNA targeting (Doepker et al., 2004; Feng et al., 2001). In fact, splicing was reported to have a protective effect on targeting by vhs (Sadek and Read, 2016). Also, SARS CoV nsp1 only degrades RNAs that are actively translated (Gaglia et al., 2012; Kamitani et al., 2009). Thus, to our knowledge, this is the first described instance of a host shutoff RNase using splicing as a targeting mechanism. The connection between splicing and PA-X degradation is evident from the reduced effect of PA-X on intronless mRNAs (Figures 4, 5) and the negative correlation between exon number and degree of degradation by PA-X (Figures 4, S3). These findings begin to shed light on the specificity of PA-X for Pol II transcripts (Khaperskyy et al., 2016). In cellular transcription, the splicing machinery associates with RNAs through interactions with Pol II, and thus only Pol II transcripts are normally spliced (Gu et al., 2013). We hypothesize that protein-protein interactions with splicing factors may bring PA-X to its transcript, an idea which is corroborated by the results of our proteomic analysis (Figure 6). Indeed the C-terminal domain of PA-X interacts with several splicing regulators (PUF60, RBM39, PRPF4) and polyadenylation proteins (CPSF5/NUDT21, CPSF6). We speculate that more exons provide more chances for PA-X to be brought to the RNA by some of these factors. In turn, this may translate into more effective turnover of RNAs with more splice sites.

A targeting strategy based on splicing offers a major benefit to the virus, because viral mRNAs are synthesized by the RdRp and most of them are not spliced. This renders them “invisible” to PA-X. That said, our published results suggest that even the two viral mRNAs that are spliced (NEP and M2) are PA-X resistant (Khaperskyy et al., 2016). However, splicing of viral mRNAs is a fundamentally different process, since the splicing machinery needs to be recruited to the RNAs separately from Pol II (Dubois et al., 2014). It is likely that viral mRNA splicing does not requires the factors that PA-X couples to, particularly since none of the potential PA-X-interacting proteins are core components of the spliceosome. The ability to easily discriminate between host and viral mRNAs may be one of the reasons IAV has evolved a targeting strategy based on splicing and not translation. For example, herpesviral mRNAs are not protected from degradation by herpesviral host shutoff RNases, presumably because there is no clear difference in their translation strategy (Abernathy et al., 2014). While herpesviruses can compensate for the degradation of their own RNAs, this self-sacrifice may not work for a virus like IAV, which has a shorter replication cycle and a simpler gene expression program.

Another conclusion of our RNAseq analysis is that PA-X with a 9-aa truncated X-ORF is essentially non-functional in the context of infection. The shutoff impairment of IAV PA(X9) is very similar to that of the PA(fs) and PA(ΔX) viruses (Fig. S1B), even though the frameshifting sequence (and thus presumably PA-X production) is not altered in the X9 virus. This finding validates the results of multiple studies using ectopic PA-X expression models (Hayashi et al., 2016; Khaperskyy et al., 2016; Oishi et al., 2015), and our PR8 results shown in Figure 1B. Previous studies indicated that at least 15 aa of the X-ORF are required for full RNA degrading activity in cells (Hayashi et al., 2016; Khaperskyy et al., 2016; Oishi et al., 2015), despite *in vitro* activity of the RNase domain alone (Bavagnoli et al., 2015; Dias et al., 2009; Yuan et al., 2009). Jagger *et al.* previously tested a premature stop mutant, and obtained a virus with an intermediate phenotype between IAV wt and PA(fs) (Jagger et al., 2012). However, they inserted the stop codon after 16 aa, which was likely not sufficient to abolish PA-X activity (Figure 1B) (Jagger et al., 2012). The finding that truncating the X-ORF is sufficient to block PA-X activity in the virus is also important, because single point mutations in the X-ORF sequence are less disruptive changes than frameshifting mutations. Thus, viruses carrying X-ORF mutations could be better tools for *in vivo* studies of PA-X function and IAV pathogenesis.

Previous studies have suggested that the positively charged residues of the X-ORF are required for nuclear localization of PA-X (Hayashi et al., 2016; Khaperskyy et al., 2016; Oishi et al., 2015). However, enforced localization of the PA-X RNase domain alone is not sufficient to rescue activity, suggesting that the X-ORF is needed for additional functions (Hayashi et al., 2016). Our BioID results corroborate this idea, since we did not only find proteins involved in nuclear import, but identified many proteins with various roles in RNA metabolism. Also, by examining the X-ORF in isolation, we likely excluded indirect interactions via RNA binding of the RNase domain, as well as interactions that are important for PA rather than PA-X function. Two proteins that we identified, nucleolin and nucleophosmin, were also reported in a proteomic study of H5N1 PA-X interactors (Li et al., 2016), suggesting they interact with at least three different PA-X variants. While nucleolin and nucleophosmin are highly abundant proteins, they are unlikely to be artifacts because they were enriched in all the X-ORF samples relative to the controls. The fact that biotinylated proteins accumulated in the nucleoli also supports the idea that nucleolin and nucleophosmin, which traffic to nucleoli, come in contact with the BirA*-fused X-ORF and full length PA-X (Figure S5C). It is unclear whether the high biotinylation in nucleoli is of functional importance, since PA-X itself is not localized specifically to this compartment (Figure S5A). Both nucleolin and nuclephosmin have multiple functions in RNA metabolism including splicing and polyadenylation, and could serve as a link between PA-X and splicing machinery (Das et al., 2013; Sagawa et al., 2011; Salvetti et al., 2016; Tarapore et al., 2006). Other hits from the BioID screen could also explain the splicing connection with PA-X. Interestingly, we did not isolate many core components of transcription or mRNA processing machinery. PA-X appears to interact mostly with factors that are considered regulators of alternative splicing (PUF60, RBM39, HNRNPK), or connect splicing with adenylation and enhance poly(A) site choice (CPSF5/NUDT21, CPSF6). It will be interesting to determine how these proteins are involved in PA-X activity. Particularly, since these proteins do not regulate the processing of all mRNAs in the cell to the same extent, interactions with these proteins could offer PA-X an additional mechanism for target discrimination. Another interesting question is whether these associations compromise the normal activity of the cellular proteins. We did not find dramatic changes in the host splicing pattern in our dataset (not shown), but we may have missed some subtle changes.

It is interesting that both PA-X and the well known influenza host shutoff factor NS1 interact with nuclear mRNA processing machinery to control gene expression (Nemeroff et al., 1998). The NS1 variants of many influenza A strains (but not PR8) cause host shutoff by binding and inhibiting a key component of the 3’-end RNA processing machinery, CPSF30 (Das et al., 2008; Nemeroff et al., 1998). It is possible that this convergence could allow the two proteins to coordinate an attack on the host. However, studies of engineered viruses and naturally-evolved viruses suggest that the activity of NS1 and PA-X may be anti-correlated to prevent cytotoxicity. For example, while the original 2009 pandemic NS1 does not bind CPSF30 and reduce host gene expression (Hale et al., 2010), currently circulating pandemic H1N1 strains carry a mutated NS1 that has increased CPSF30 binding and host shutoff activity (Clark et al., 2017; Nogales et al., 2018). However, these same strains have also accumulated PA-X mutations that reduce PA-X activity, suggesting that having two highly active host shutoff proteins may impair viral fitness (Nogales et al., 2018). In general, host mRNA processing may be a hub of regulation for influenza, since it can easily allow for discrimination of host RNAs that are processed *vs.* viral mRNAs that are not. At the same time, interactions with host processing do not affect Pol II activity, which is important given that Pol II transcription is required for viral replication (Lamb and Choppin, 1977). It will be interesting in the future to observe how PA-X and NS1 interact in viruses that, unlike PR8, use NS1 to block mRNA processing.

Collectively, our results affirm the importance of PA-X for the viral replication cycle, as they show that the ability of the virus to regulate host gene expression is severely reduced in the absence of PA-X. Moreover, we have uncovered a unique mechanism of host RNA targeting that can allow PA-X to not only distinguish between host and viral targets, but also among cellular targets. For example, the intronless mRNA TAF7, which we examined in Figures 2B, 2C, is a component of Pol II pre-initiation complexes. Therefore, the selectivity of PA-X could have repercussions for the viral replication cycle. By continuing to define the PA-X mechanism of action, we will gain important insights into how host shutoff allows the virus to usurp host biosynthetic machinery, and expand our knowledge of the link between PA-X and IAV pathogenesis.

## EXPERIMENTAL PROCEDURES

### Plasmids

pCR3.1-PA-X-myc (with PR8 PA-X) (Khaperskyy et al., 2014, 2016), pCR3.1-PA-X_TN/CA/7-myc (Khaperskyy et al., 2016), pd2GFP-HR (Lee and Glaunsinger, 2009) were previously described. The luciferase constructs with and without the intron were a kind gift from Gideon Dreyfuss (Younis et al., 2010). pHW-PA(X9) and pHW-PA(∆X) were generated from pHW-193 (kind gift from R. Webby) and pHW-PA(fs) vectors (Khaperskyy et al., 2016), respectively, using Phusion site-directed PCR mutagenesis to introduce the TAG stop codon in +1 ORF (synonymous ATT to ATA substitution at PA Ile-201 codon, TTG to TAG substitution at PA-X Leu-201). pCR3.1-PA-X_Udorn-myc was generated by PCR amplifying the 5’ portion of the segment 3 RNA from a PolI-Udorn construct (kind gift from A. Mehle), adding a single nucleotide deletion to shift the frame of the X-ORF, and inserting into the SalI-MluI sites of a pCR3.1-C-terminal-myc backbone using Gibson cloning (HiFi assembly mix, New England Biolabs). pTRIPZ_PA-X_Udorn-myc was generated by PCR amplifying PA-X-Udorn-myc from pCR3.1-PA-X_Udorn-myc, and inserting it into the backbone of pTRIPZ-RFP_SV40_3’UTR (Khaperskyy et al., 2016) after RFP excision with AgeI and ClaI using Gibson cloning (HiFi assembly mix, New England Biolabs). pCMV-IFNL2 cDNA, genomic and single intron constructs were generated by PCR amplifying the full human IFNL2 cDNA, the genomic locus or combinations of fragments of the two, and inserting into the pd2eGFP-N1 construct (Clontech) after the GFP was excised using NheI and NotI. HiFi assembly mix (New England Biolabs) was used to make these constructs. Expression vector for the biotin ligase from *E. coli* with the R118G mutation BirA* (pcDNA3.1-myc-BioID2-MCS) (Roux et al., 2012) was obtained from Addgene (#74223) and the BirA* ORF was amplified by PCR and inserted into the pCR3.1-myc vector (Khaperskyy et al., 2012) between KpnI and EcoRI sites to generate pCR3.1-BirA*-myc. The PCR-amplified X-ORF sequences from pCR3.1-PA-X-myc and pCR3.1-PA-X(4A)-myc (Khaperskyy et al., 2016) were inserted in frame with BirA* ORF using EcoRI and MluI to generate pCR3.1-BirA*-X61-myc and pCR3.1-BirA*-X61(4A)-myc, respectively. X-ORF coding sequence from A/California/7/2009(H1N1) strain was amplified from pHW-C3 vector (Slaine et al., 2018) and inserted in frame with BirA* ORF using EcoRI and XhoI to generate pCR3.1-BirA*-X41(CA/7) vector.

### Cell lines, lentiviral transduction and transfections

HEK293A, HEK293T, A549 cells and derivatives were cultured in Dulbecco’s modified Eagle’s medium (DMEM) supplemented with 10% fetal bovine serum (Hyclone) at 37°C and 5% CO_2_. A549-iPA-X_PR8 and A549-iPA-X-D108A_PR8 were previously described (Khaperskyy et al., 2016). A549-iPA-X_Udorn were generated by transducing A549 cells (ATCC) with lentiviruses containing pTRIPZ-PA-X_Udorn-myc. Lentiviral packaging was carried out using the packaging plasmids psPAX2 and pMD2 (Addgene #12260, #12259). For experiments using iPA-X cells, cells were treated with 0.2 μg/ml doxycycline for 18 hrs to induce PA-X expression prior to RNA sample collection. For the RNAseq experiments, untransduced A549 cells were also treated with doxycycline to serve as the control. For experiments using IFN-λ2 constructs and the β-globin reporter, HEK293T cells were plated in 24-well or 6-well plates (for fractionation experiments) and transfected with 800 ng/ml total DNA (including 50 ng/ml PA-X construct) using polyethylenimine (PEI). Cells were collected 24 h later for fractionation and/or RNA extraction and purification.

### Viruses and infections

Wild-type influenza A virus A/Puerto Rico/8/1934 H1N1 (PR8) and the mutant recombinant viruses PR8 PA(X9), PR8 PA(fs) and PR8 PA(∆X) were generated using the 8-plasmid reverse genetic system (Hoffmann et al., 2000) as previously described (Khaperskyy et al., 2012). Viral stocks were produced in MDCK cells and infectious titers determined by plaque assays in MDCK cells using 1.2% avicel overlays as described in Matrosovich *et al*. (Matrosovich et al., 2006). A549 cell monolayers were mock-infected or infected with the wild-type or mutant viruses at MOI = 1 for 1 h at 37°C. Then monolayers were washed briefly with PBS, fresh infection media (0.5% BSA in DMEM supplemented with 20 μM L-glutamine) was added and cells incubated at 37°C in 5% CO_2_ atmosphere for 12 or 15 h prior to RNA isolation or preparation of lysates for western blotting. For immunofluorescence microscopy analysis cells grown on glass coverslips were infected as described above and fixed at 15 h post-infection using 4% paraformaldehyde in PBS.

### Preparation of cell lysates containing biotinylated proteins

HEK 293T cells grown on 10-cm dishes were washed briefly were transfected with BirA* fusion protein expression constructs using PEI. 6 hours post transfection media was changed to 10% FBS DMEM supplemented with 50 μM biotin (Sigma). 24 hours post-transfection (18 h post-biotin addition) cells were washed and collected in ice cold PBS, and centrifuged at 250 x *g* for 5 minutes at 4°C. Cell pellets were resuspended in 500 μl RIPA buffer (50 mM Tris-HCl pH 7.4, 150 mM NaCl, 1% Igepal, 0.5% sodium deoxycholate, 0.1% SDS) with protease inhibitor cocktail (P8340, Sigma) and lysed at 4°C for 1 hour with gentle agitation, followed by passing through a 21-gauge needle. Lysates were cleared by centrifugation at 4°C for 20 min at 20,000 x *g*.

### Neutravidin pull-down

60 μl of 50% slurry of High Capacity Neutravidin Agarose Beads (Thermo) was used for each 500 μl of clarified whole cell lysate. Beads were equilibrated in RIPA buffer by washing three times for 10 mins at 4°C. In one of the BirA*-X61 experimental runs, 1 μl of 500x RNase A (100 μg, Qiagen) was added to each sample to remove non-specific interactors. The lysate was then incubated for 5 minutes at room temperature before loading onto the beads. Untreated samples were loaded directly onto the beads post washing. 1 mg of protein sample was loaded to beads in 1.5 mL Eppendorf tubes, which were then placed on a rotator overnight at 4°C, and collect with centrifugation at 400 x g for 1 minute at 4°C. Beads were washed with RIPA buffer three times, followed by three washes with TAP buffer (50 mM HEPES-KOH pH 8.0, 100 mM KCl and 10% glycerol).

### Mass spectrometry sample preparation

Beads were resuspended in 50 mM triethylammonium bicarbonate (TEAB) buffer (Sigma). 24 mM DTT and 32 mM IAcNH2 were added sequentially. Beads were then incubated for 30 minutes at 37°C, washed with 50 mM TEAB and centrifuged for 1 min at 400 x *g* before resuspending in 50 mM TEAB. On-bead trypsin (PierceTM Trypsin protease, MS-Grade; Thermo Scientific) digest was performed with 1 μg trypsin in 50 mM TEAB buffer, shaken overnight at 37°C. Samples were acidified with 1 μl trifluoroacetic acid (TFA) and 3 μl Formic acid until a pH lower than 3 was achieved. Trypsinized peptides were collected by puncturing a hole in the bottom of the 1.5 mL Eppendorf tube using a 30-gauge needle, placing it in a 2 mL Eppendorf tube and spinning it at 2000 rpm for 1 minute at room temperature. Beads were washed with 50 mM TEAB prior to desalting. Samples were desalted with Oasis/SepPak Desalting columns, eluted sequentially in 1 mL 50% ACN/0.1% TFA and 500 μl 70% ACN/0.1% TFA. Combined eluted samples were dried in a Thermo SPDIIIV speed vacuum centrifuge and frozen at −20°C.

### Reductive dimethylation and quantitative mass spectrometry

Quantitative mass spectrometry analysis via reductive dimethylation enabled measurements of relative abundance of biotinylated proteins in each experimental condition (Hsu et al., 2003). In reductive dimethylation, formaldehyde molecules with different combinations of stable hydrogen and carbon isotopes are conjugated to peptide samples. Dried protein samples were resuspended by sonication for 15 minutes in 50 mM TEAB. BirA*, BirA*-XORF (X61 or X41) and BirA*- X61(4A) samples were labeled with light, medium and heavy isotopes respectively. 8 μl formaldehyde (Sigma) were added to the light, 15 μl D2-formaldehyde (Cambridge Isotope Laboratories Inc.) to the medium and 15 μl D2-C13-formaldehyde (Aldrich) to the heavy samples. Reactions were incubated for 5 minutes at room temperature. Once incubation was completed, 0.51 M NaCNBH3 (sodium cyanoborohydride; Fluka) was added to the light and medium samples, while 0.51 M NaCNBD3 (sodium cyanoborodeuteride; Aldrich) was added to the heavy reaction, to label terminal amines. All three reactions were incubated for 1 hour at room temperature before being combined into a single tube at a 1:1:1 ratio. The combined sample was acidified, desalted and dried as described above. Samples were resuspended in 3% ACN/0.1% formic acid and sonicated for 15 min to prepare for mass spectrometry. Mass spectrometry and peptide identification was performed at Dalhousie Proteomics CORE Facility by Dr. Alejandro Cohen (https://medicine.dal.ca/research-dal-med/facilities/proteomics.html). Proteome Discoverer software (Thermo) was used for protein identification. Functional Protein Association Network analysis was conducted on 29 selected protein hits using online STRING version 10.5 (https://string-db.org/) (Szklarczyk et al., 2017).

### Cell fractionation

Fractionation was performed as described previously (Gagnon et al., 2014) with a few modifications. Briefly, cells were collected 24 h after transfection, centrifuged at 500 x *g* for 5 min at 4°C, then washed with PBS and counted. Equal numbers of cells were aliquoted into two tubes, one for the whole cell lysates collection and one for fractionation. Cells were pelleted again, and lysed on ice for 10 mins in 250 μl ice-cold hypotonic lysis buffer (10 mM Tris pH 7.5, 10 mM NaCl, 3 mM MgCl_2_, 0.3% (vol/vol) NP-40, 10% (vol/vol) glycerol in nuclease-free water) supplemented with 100 U RNasin (Promega). For the whole cell lysate, Trizol (Life Technologies) was added directly to the lysate to extract RNA. For nuclear/cytoplasmic fraction, the lysate was centrifugated at 1000 x *g* for 3 min at 4°C to pellet membrane and nuclei. The supernatant was collected as the cytoplasmic fraction and Trizol was added to it to extract RNA. Finally, the nuclear pellet was washed 3 times with 1 ml hypotonic lysis buffer and collected by centrifugation at 200 x *g* for 2 mins at 4°C, then lysed directly in Trizol to extract nuclear RNA.

### RNA purification, cDNA preparation and qPCR

For fractionation experiments, RNA was purified using Trizol. 1 ml Trizol, 2 μl glycogen and 200 μl chloroform (Fisher Scientific) were added to each fraction. Samples were then centrifuged at 16,000 x *g* for 15 min at 4°C, and the aqueous layer was collected. RNA was precipitated by addition of 700 μl isopropanol and incubation for 10 min at room temperature, followed by centrifugation at 16,000 x *g*, 4°C for 20 min. The pellet was washed with 75% ethanol, and resuspended in RNase-free water. For other transfection experiments, RNA was extracted from cells and purified using the Quick-RNA miniprep kit (Zymo Research), following manufacturer’s protocol. In all cases, the RNA was treated with Turbo DNase (Life Technologies), then reverse transcribed using iScript supermix (Bio-Rad) per manufacturer’s protocol. In the fractionation experiment, the same cell equivalents of total, nuclear and cytoplasmic fraction were used for these steps. qPCR was performed using iTaq Universal SYBR Green supermix (Bio-Rad), on the Bio-Rad CFX Connect Real-Time System qPCR and analyzed with Bio-Rad CFX Manager 3.1 program. The primers used are listed below.

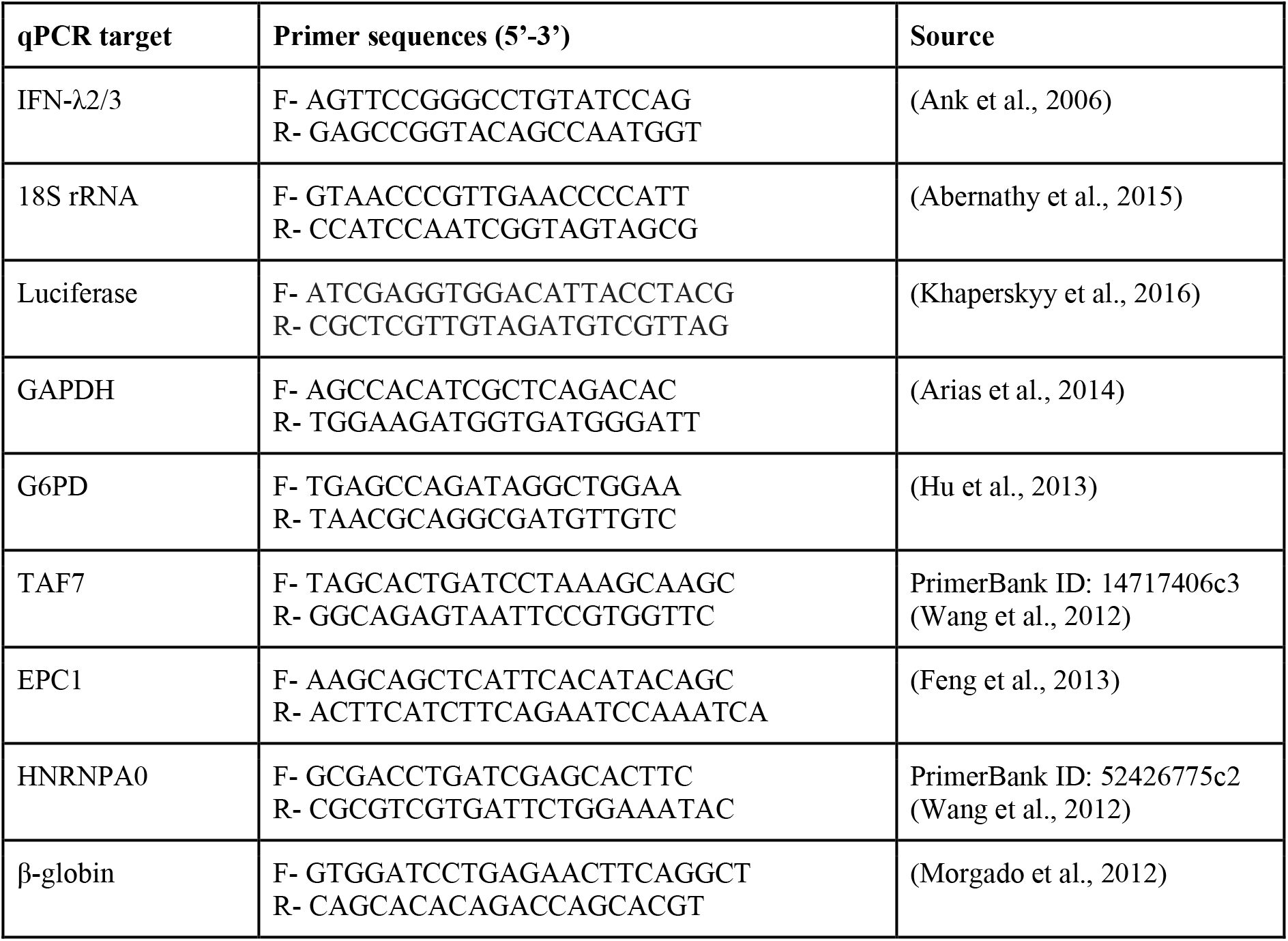

### Intron splicing verification

Proper splicing of each intron for all IFN-λ2 construct was verified by PCR amplification across each splice sites using primers listed below. PCR products were run on a 2% agarose gel containing HydraGreen safe DNA dye (ACTGene) and imaged with a Syngene G:Box Chemi XT4 gel doc system.

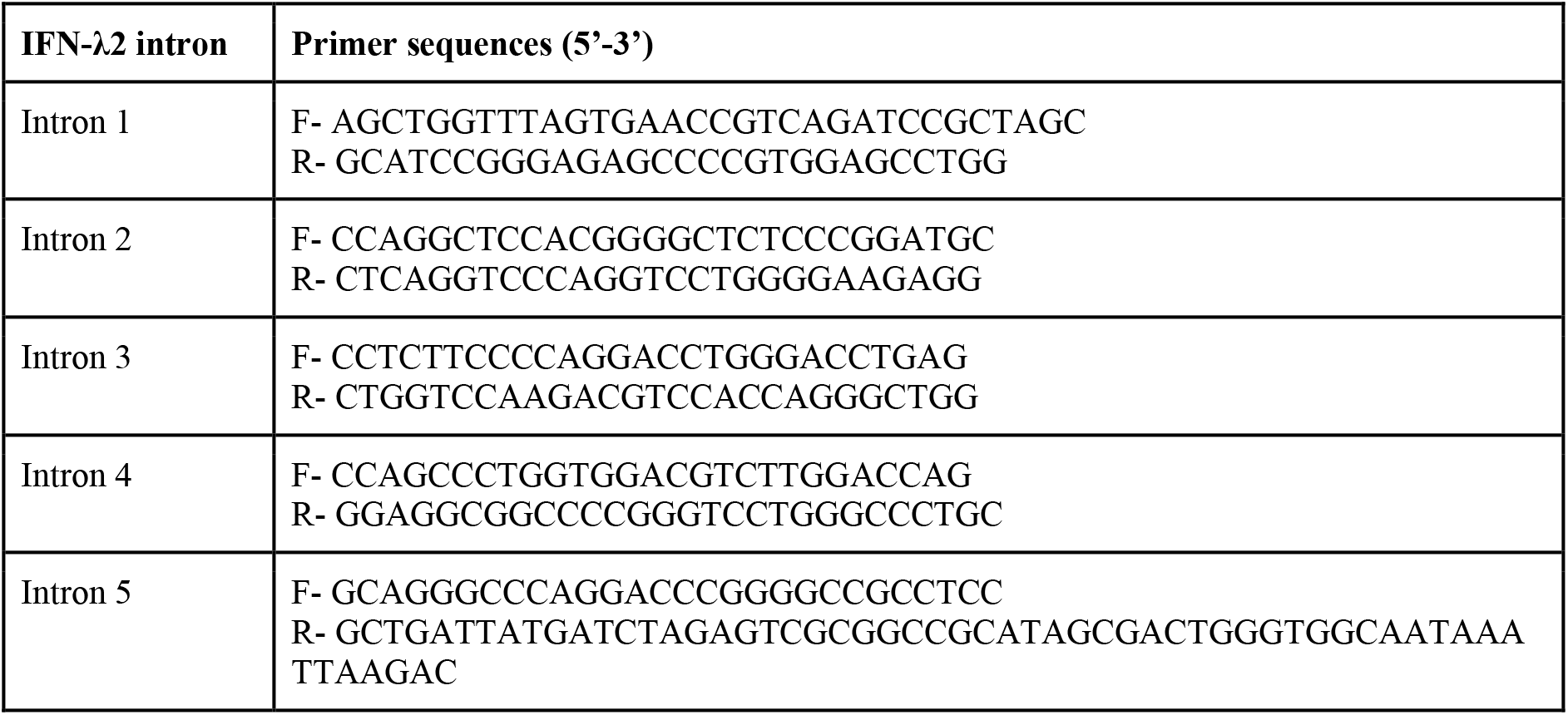

### Statistical analysis

For Figures 1B, 2C, 2D, 5, statistical analysis and plotting were done in GraphPad Prism v7.0d software using the test recommended by the software and indicated in the figure legends. Generally, ANOVA followed by a corrected pairwise test (Tukey’s or Dunnett’s) was used when more than two samples were analyzed and Student’s *t* test when two samples were compared. For Figures 1C-D, 2A-B, 4, statistical analysis and plotting were done using Python2.7 and the NumPy, SciPy and matplotlib libraries. The Kolmogorov-Smirnoff test was used to compare populations and Spearman’s correlation coefficient to analyze the relationship between certain variables and RNA down-regulation.

### RNA-seq

For RNAseq analysis of PA-X-expressing cells, A549 and A549 iPA-X cells were induced with 0.2 μg/ml doxycycline for 18 hours. For RNAseq analysis of infected cells, A549 cells were infected with PR8 wt, PA(fs), PA(X9), and PA(ΔX) viruses for 15 hours. RNA lysates were collected and purified using the RNeasy kit (Qiagen). 650-750 ng of RNA were mixed with ERCC ExFold RNA spike-in mix (0.65 μl or 0.75 μl, respectively, Invitrogen/ThermoFisher) prior to the start of library preparation. The spike-in controls were included to better normalize the final RNA levels. This particular kit contains two mixes that can be used for the control *vs.* test samples and that have known fold change differences. This information can be used to re-calibrate the samples. Prior to library preparation, the levels of select human mRNAs were tested by RT-qPCR, to confirm that PA-X overexpression/IAV infection had the expected effect. Libraries were prepared using the TruSeq Stranded Total RNA Library Prep Kit with Ribo-Zero Human (Illumina) following the manufacturer’s protocol. Library preparation was evaluated using a Fragment Analyzer (Advanced Analytical Technologies, Inc.) at the Tufts University Core Facility - Genomics Core. High-throughput sequencing was carried out by the Tufts Genomics facility on a HiSeq 2500. Single-end 50 nucleotide reads were obtained with a multiplexing strategies, using a total of four total lanes. For the replicates, the spike mixes for the samples and the barcoded primers were switched, in order to control for potential biases.

### Read alignment and bioinformatic analysis

Reads were aligned with Tophat2 v2.1.1 (Kim et al., 2013) to hg19, IAV PR8 and the ERCC spike sequences. Default settings were used, except--library-type fr-firststrand, and a gtf of the hg19 annotation was provided as reference. Table S1 summarizes the results of the alignment. FPKMs were computed using Cufflinks v2.2.1 (Trapnell et al., 2012). Default settings were used, except--library-type fr-firststrand and −u, and a gtf of the hg19 annotation was provided as reference. Only previously annotated RNAs with a level of >1 FPKM in all the control samples (A549 + dox or mock-infected A549) were used in further analyses. The FPKMs for these RNAs were converted to attomoles of RNA based on the known concentration of the spike-in controls. Table S2 shows the high correlation in measured RNA levels between replicate samples. The absolute RNA levels in the replicates were averaged, and the relative RNA levels (RNA ratio) in infected *vs.* mock infected or PA-X-expressing *vs.* control cells was computed. All downstream analysis was carried out on the relative levels (ratio) in log2 scale. Tables S3 and S4 summarize the results of the RNAseq analyses. RNAseq data is available on the GEO database, entry GSE120183 (raw read data and processed data included here as Tables S3 and S4). The hg19 annotation was used to derive the number of exons, length of transcripts and GC content for the analyses in Figures 4, S3. For analysis of intronless RNAs, only RNAs that were longer than 300 nt were used, to exclude small RNAs that are transcribed by Pol III or are produced through processing of longer transcripts. The length distribution of the remaining intronless RNAs was similar to that of the spliced RNAs. All RNAs were used for the analysis of PA-X down-regulation *vs.* length, exon number, and GC content. *k*-means clustering analysis was carried out using the Cluster 3.0 program (http://bonsai.hgc.jp/~mdehoon/software/cluster/software.htm). All eight datasets were used to generate the clusters, even though only the mRNA levels for select samples are plotted in Figures 3A-C. Because in *k*-means clustering the number of clusters is user-defined, clustering was attempted with three, four, and five clusters. Four clusters were chosen, because they provided more granularity. For example, they identified a group of RNAs that were only PA-X-dependent in the ectopic expression system. Initializing the program with more than four clusters led to separation of PA-X targets in multiple groups different only by the extent of down-regulation, but did not identify other patterns of gene expression. Gene ontology (GO) term analysis was carried out on the DAVID server. Other analyses were done using custom scripts in Python2.7. For the ISG analysis (Figure 3E) the list of ISG tested by Schoggins *et al.* was used (Schoggins et al., 2011).

### Western blotting and immunofluorescence

Western blotting and immunofluorescence were carried out as previously described (Khaperskyy et al., 2014, 2016). For detection of biotinylated proteins HRP-conjugated streptavidin (3999, Cell Signaling) and Alexa-Fluor-488-conjugated streptavidin (Molecular Probes) were used. Antibodies: mouse monoclonal antibody to PABP1 (sc-32318, Santa Cruz Biotechnology); goat polyclonal antibody to influenza virus (ab20841, Abcam); rabbit polyclonal to nucleolin (ab22758, Abcam).

## AUTHOR CONTRIBUTIONS

Conceptualization, C.M., D.A.K., M.M.G.; Investigation, L.G., B.K.P., S.K.S, Y.K., D.A.K., M.M.G.; Formal analysis, C.H.R., M.M.G.; Writing – original draft, L.G., C.M., D.A.K., M.M.G.; Funding acquisition, C.M., M.M.G.

## ACKNOWLEDGMENTS

We thank Albert Tai and the personnel of the Tufts University Core Facility - Genomics Core for help with the RNAseq. We thank Alejandro Cohen of the Dalhousie Proteomics and Mass Spectrometry Core Facility for support of BioID proteomics analysis. We thank Andrew Mehle and Richard Webby for constructs, Claire Moore and Andrew Bohm for suggestions and feedback, and members of the Gaglia and McCormick labs for critical reading of the manuscript. This work was supported by National Institutes of Health grant R01 AI137358 (to MMG) and Canada Institutes for Health Research grant MOP-136817 (to CM). LG is supported by National Institutes of Health training grant T32 GM007310.

## SUPPLEMENTAL INFORMATION

**Table S1.**
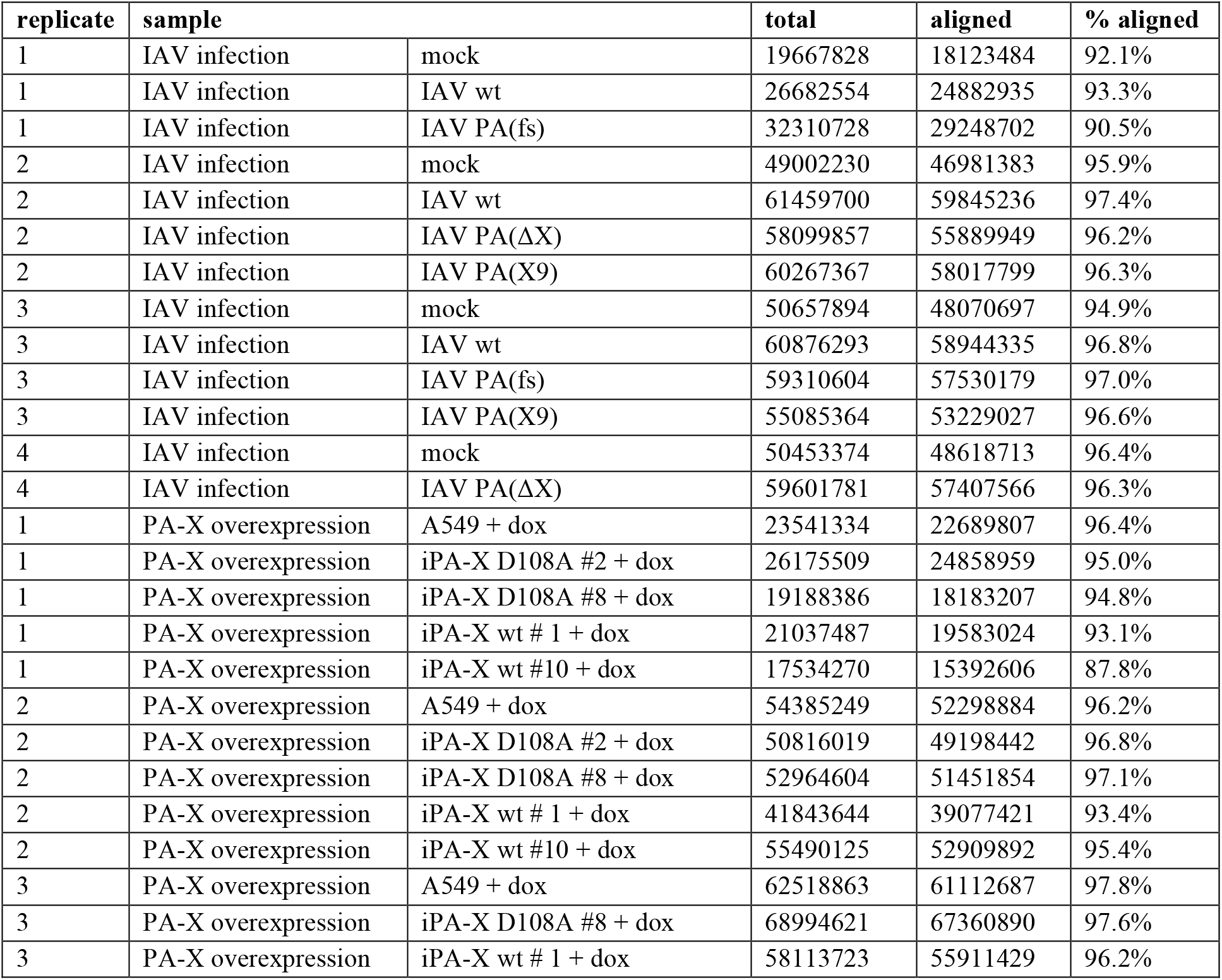
Summary of RNAseq read data. Number of total and aligned reads per sample.

**Table S2:**
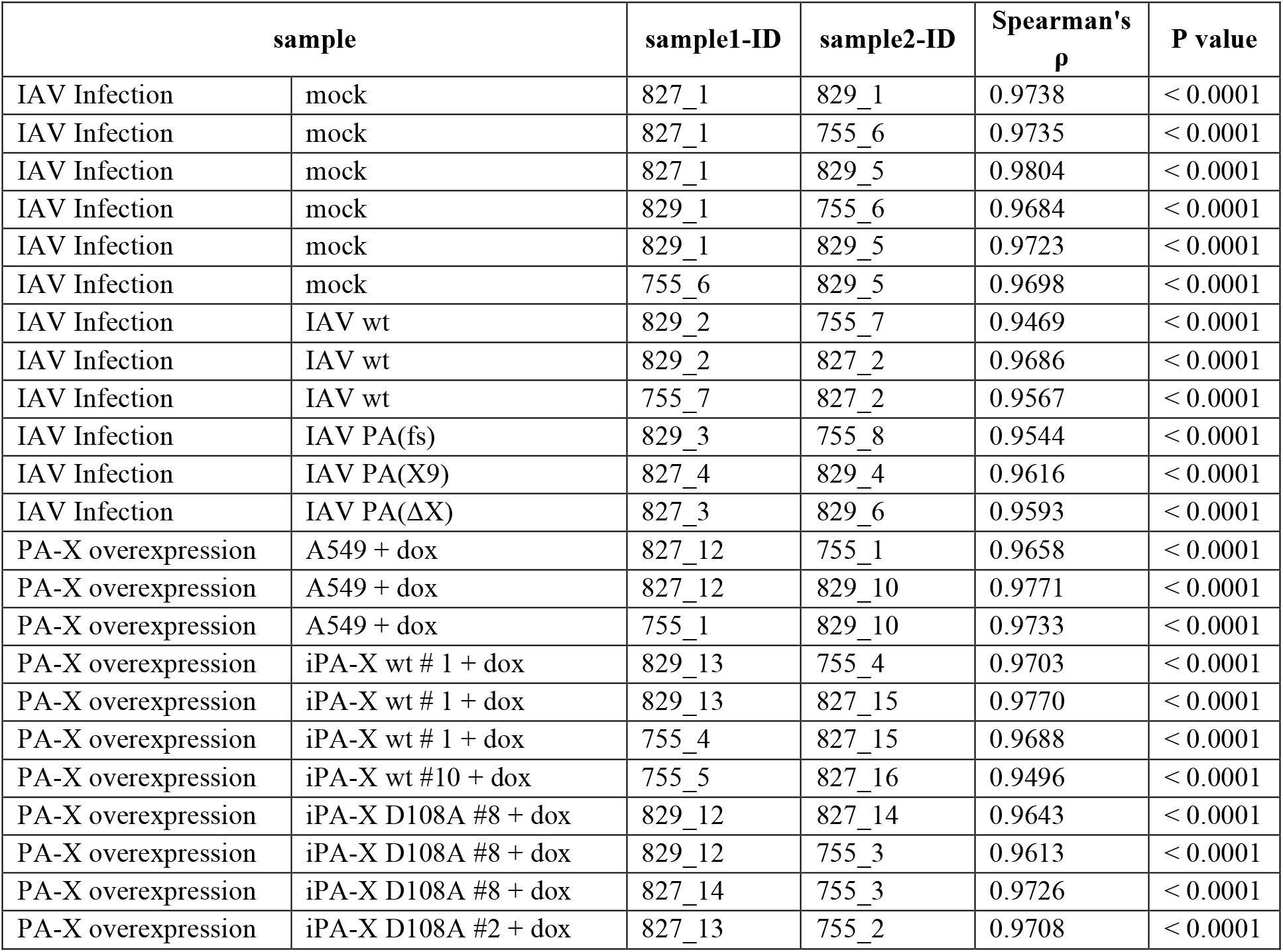
Correlation between RNA levels measurements in RNAseq replicates. List of correlation coefficients from pairwise comparison of replicates of theRNA seq (Spearman’s ρ).

**Table S3 (.txt file): Relative RNA levels in IAV-infected cells**

Summary of relative RNA levels in IAV-infected *vs.* mock infected cells for all four IAV strains used. Values are in log2 attomoles. Fold-change from replicate averages and standard deviations are reported.

**Table S4 (.txt file): Relative RNA levels in PA-X-expressing cells**

Summary of relative RNA levels in PA-X-expressing *vs.* controls cells for all four iPA-X cell lines used. Values are in log2 attomoles. Fold-change from replicate averages and standard deviations are reported.

**Table S5 (xlsx file): Summary of BioID results.**

List of proteins identified in each of the experimental runs and of the 29 high-confidence hits.

**Figure S1 (related to Figure 1).**
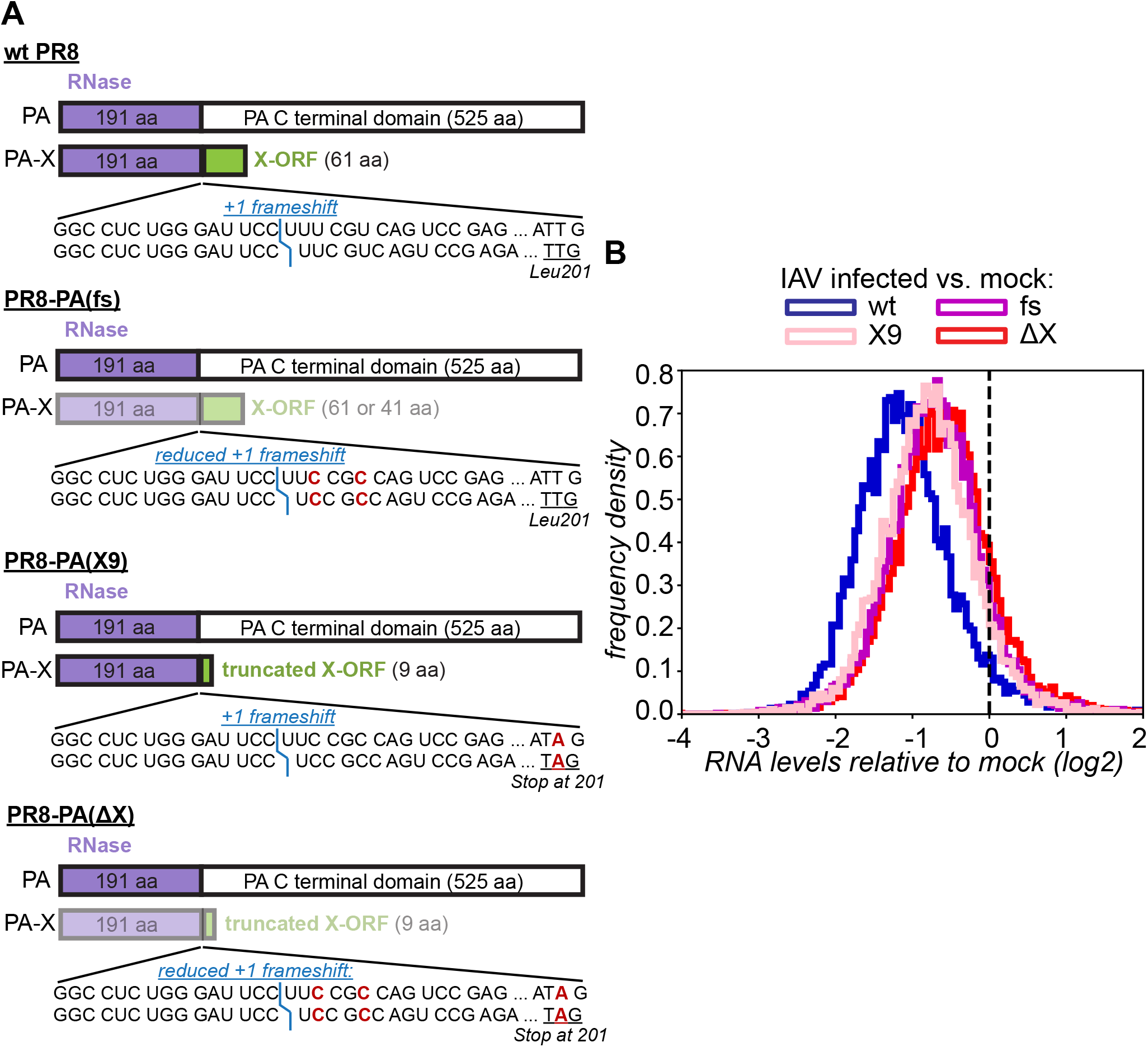
Reduction in frameshifting and truncation of the X-ORF have similar effects on host shutoff during infection. (A) Diagrams of mutations in the PR8 PA(fs), PA(X9), PA(ΔX) viruses. The less intense colors represent lower levels of the PA-X protein. The position of the frameshift is marked in blue. The mutated nucleotides in the frameshifting sequence and at PA-X codon 201 are marked in red. (B) RNA was collected from infected A549 cells 15 hrs after infection with wt PR8 or the PR8 PA-X mutant viruses (PA(fs), PA(X9), PA(ΔX)), and the levels of all RNAs were measured by RNA-seq. The ratio between the levels in IAV-infected *vs.* mock-infected cells was computed for each RNA and the distribution of the ratios is plotted as a frequency histogram (wt and PA(ΔX) plots are the same as Figure 1C). N ≥ 2.

**Figure S2 (related to Figure 1).**
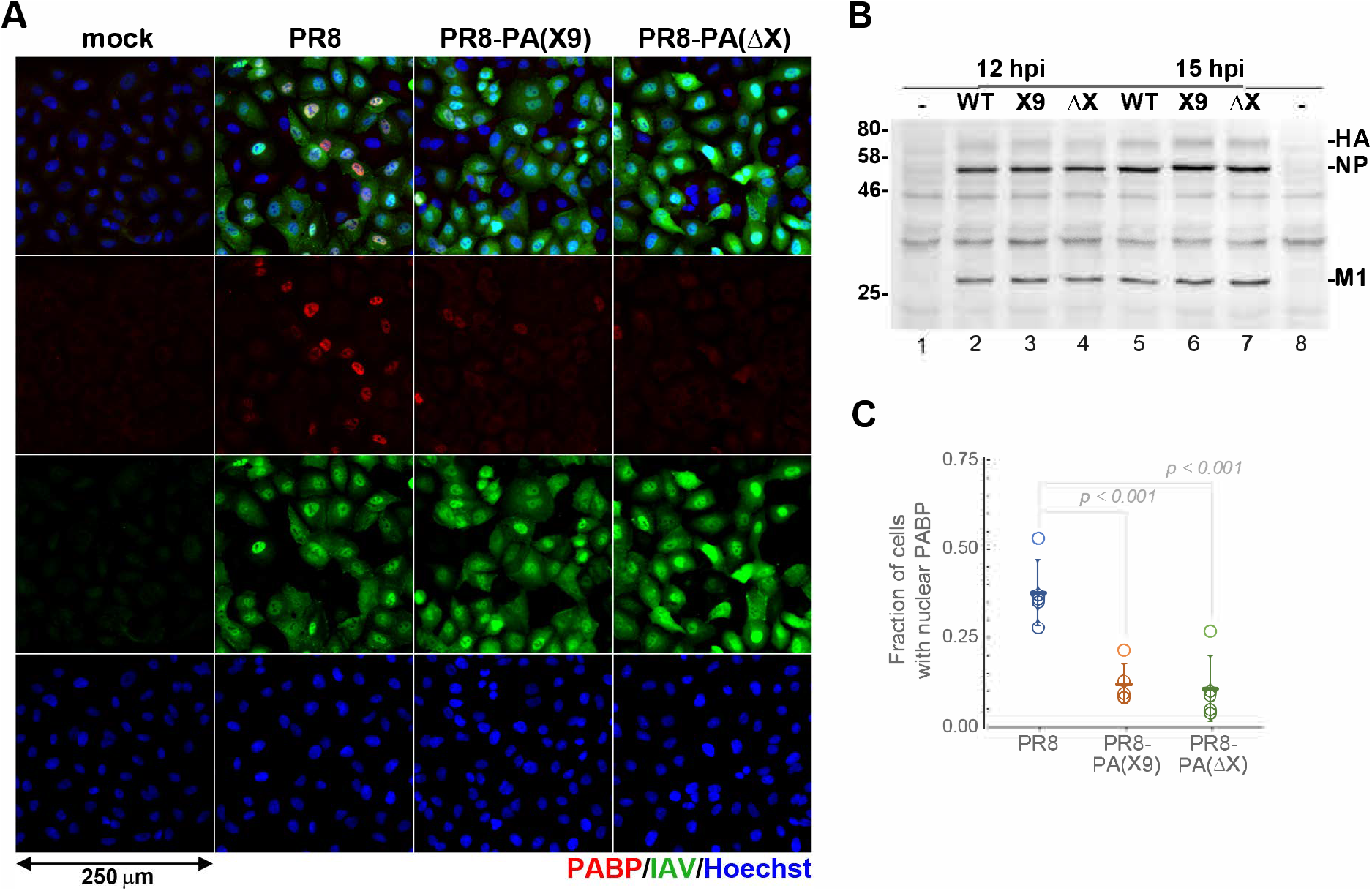
Infection rates are similar, but nuclear accumulation of PABP is reduced in cells infected with PA-X mutant viruses. A549 cells were infected with PR8 wt, PR8-PA(X9), or PR8-PA(ΔX) at MOI = 1. (A) Immunofluorescent staining of virus-infected cells at 15 h post infection (hpi) using antibodies to IAV proteins (green) and PABP1 (red). Nuclei are visualized with Hoechst dye (blue). (B) Western blot analysis of whole cell lysates collected at 12 and 15 hpi showing viral protein accumulation in infected cells. Viral HA, NP, and M1 proteins were detected using the same polyclonal anti-IAV antibody used in A. (C) Nuclear accumulation of PABP at 15 hpi was quantified from A. The fraction of infected cells with nuclear PABP staining is plotted. Each open circle represents an independent infection experiments, in which > 200 cells from at least 3 random fields of view were counted. Horizontal lines show average values and error bars correspond to standard deviations (N = 5). *p* values were determined using unpaired *t* test.

**Figure S3 (related to Figure 4).**
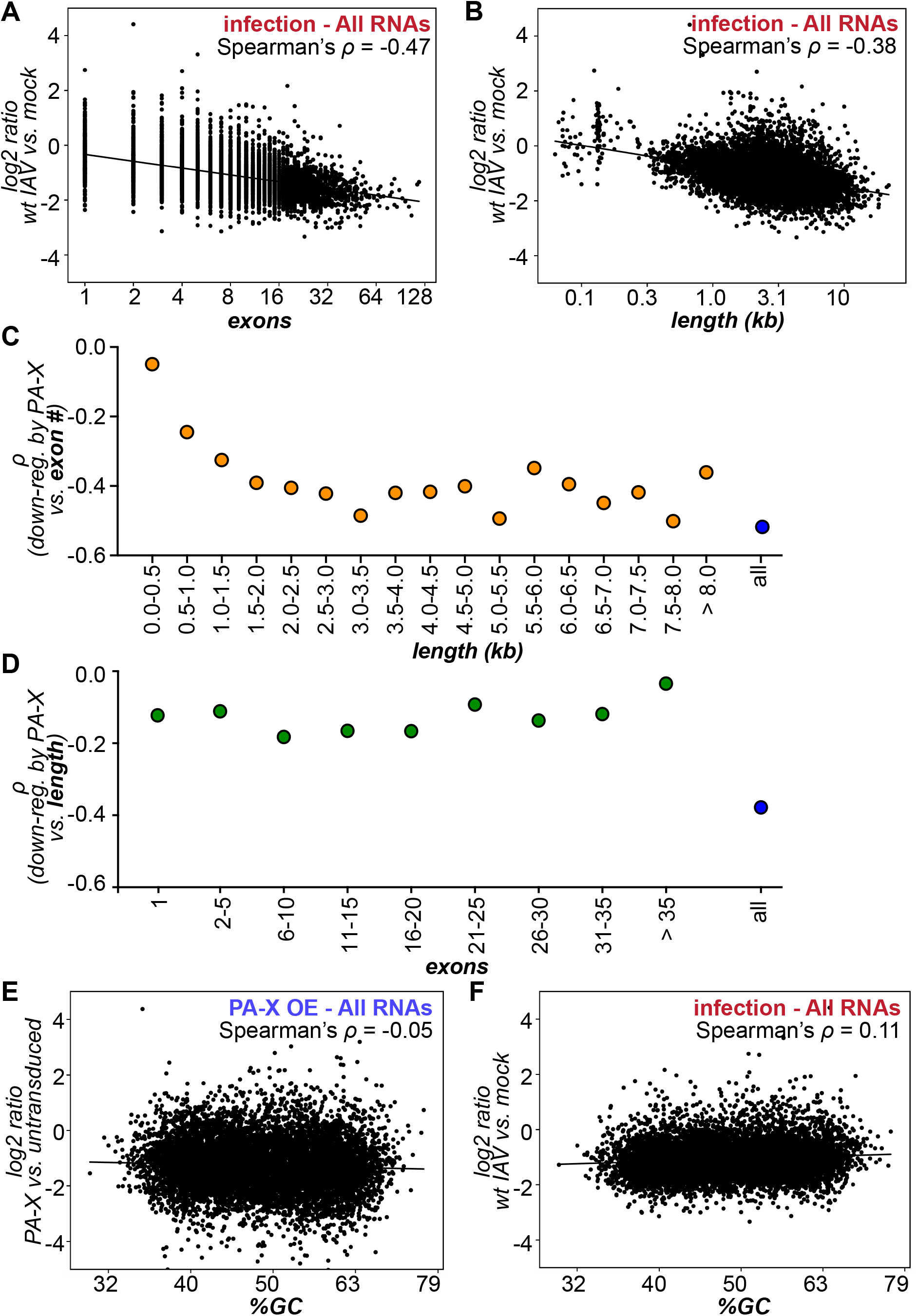
Degradation by PA-X correlates with number of exons and mRNA length, but not GC content of the mRNA. (A-B) The relative RNA levels in wt IAV infected *vs.* mock infected cells are plotted against the number of exons (A, log2 scale) or transcript length in kb (B, log10 scale). (C-D) The Spearman’s correlation coefficient (ρ) between the relative RNA levels in PA-X-overexpressing *vs.* control cells and the number of exons (C) or the RNA length (D) is plotted for groups of RNAs of the indicated length/exon number interval. Each data point represents between 99-2626 RNAs. The ρ for the entire populations are plotted in blue for comparison. (E-F) The relative RNA levels in PA-X overexpressing *vs.* control cells (E) or wt IAV infected *vs.* mock infected cells (F) are plotted against the percentage of GC of the transcript.

**Figure S4 (related to Figure 5).**
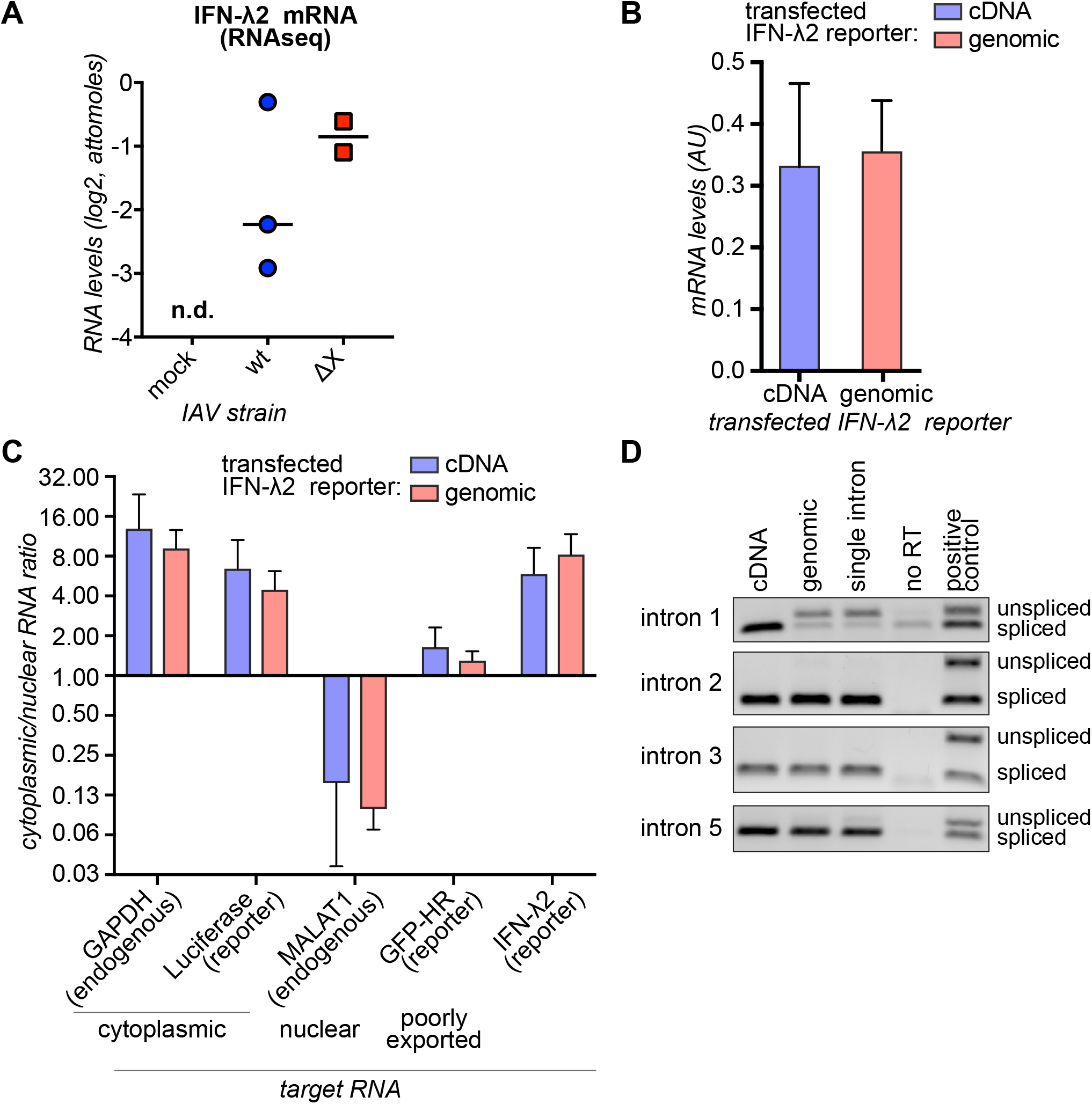
IFN-λ2 mRNA is expressed and exported at similar levels from both a genomic and a cDNA reporter constructs. (A) The attomoles of IFN-λ2 mRNA in cells infected with wt IAV or IAV PA(ΔX) are plotted. Each point represents a biological replicate. The IFN-λ2 mRNA was undetectable in mock-infected cells (n.d. = not detected). N ≥ 2 (B) HEK293T cells were transfected with reporters expressing IFN-λ2 mRNA from cDNA or the genomic locus. RNA samples were collected 24 h after transfection and analyzed by RT-qPCR. The levels of IFN-λ2 in the absence of PA-X are plotted relative to 18S cellular rRNA. All values represent mean ± standard deviation. N = 4. (C) HEK293T cells were transfected with reporters expressing a luciferase mRNA, a GFP mRNA ending in a hammerhead ribozyme (GFP-HR) and IFN-λ2 mRNAs from cDNA or the genomic locus. Cells were fractionated into nuclear and cytoplasmic fraction, and the ratio between cytoplasmic and nuclear RNA levels was calculated after normalization to 18S rRNA. The cytoplasmic/nuclear ratio for the well-exported endogenous GAPDH mRNA and transfected luciferase mRNA, the nuclear-retained endogenous MALAT1 non-coding RNA, the poorly exported transfected GFP-HR mRNA and the test IFN-λ2 mRNAs are plotted. All values represent mean ± standard deviation. N = 4. (D) The cDNAs from vector-transfected cells from Figure 5C were PCR amplified across the different introns to confirm splicing. The amplified PCR products are shown (gel image is representative of four experiments). A 1:1 mix of the constructs for IFN-λ2 cDNA and genomic serves as a control to check that both products could be simultaneously amplified.

**Figure S5 (related to Figure 6).**
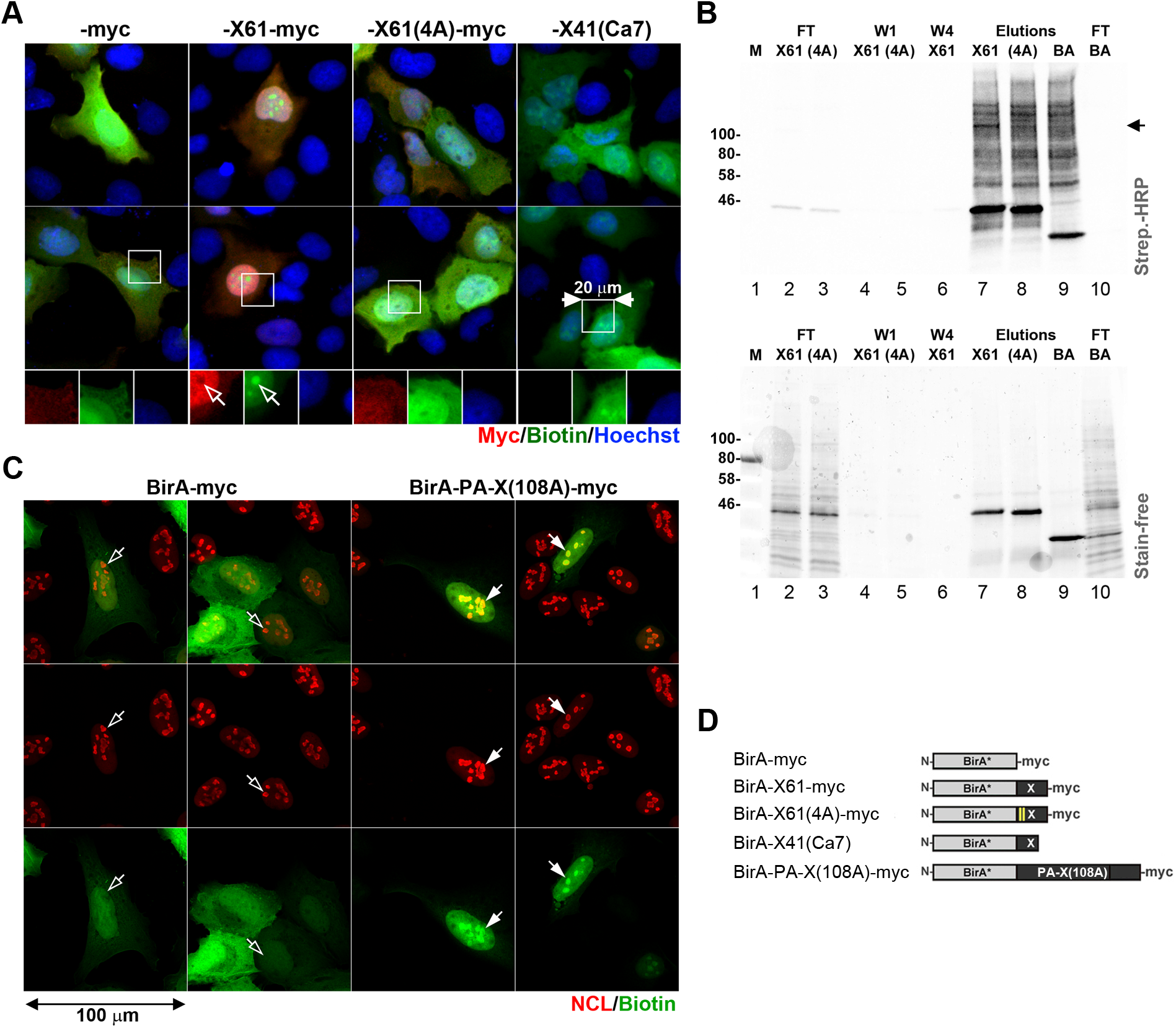
BirA*-X-ORF mediates biotinylation of cellular proteins. (A) Immunofluorescent staining of biotinylated proteins in HEK293A cells transfected with BirA*-myc (-myc), BirA*-X61-myc (-X61-myc), BirA*-X61(4A)-myc (-X61(4A)-myc), or BirA*-X41(Ca7) expression constructs. The localization of myc-tagged fusion proteins was visualized with anti-myc antibody (red), biotinylated proteins were stained with Streptavidin-Alexa-Fluor-488 conjugate (Biotin, green), and nuclei were stained with Hoechst dye (blue). Open arrows indicate strong nucleolar biotin staining in BirA*-X61-myc expressing cells. (B) Flow through (FT), wash 1 (W1), wash 4 (W4), and elution fractions of neutravidin agarose pulldown of proteins biotinylated using BirA*-X61-myc (X61), BirA*-X61(4A)-myc (4A), or BirA*-myc (BA) were analyzed by western blot using Streptavidin-HRP conjugate (top panel). Total protein in the same fractions was detected using Bio-Rad Stain-free chemiluminescent reagent (bottom panel). Arrow indicates position of a major band corresponding to the size of nucleolin (apparent molecular weight ~110 kDa) that is enriched in BirA*-X61-myc biotinylated protein mix compared to proteins biotinylated by the other two baits. (C) Immunofluorescent staining of biotinylated proteins in cells transfected with BirA*-myc or the BirA* fused to full-length catalytically inactive PA-X mutant, BirA*-PA-X(108A)-myc. Biotinylated proteins were stained with Streptavidin-Alexa-Fluor-488 conjugate (Biotin, green), and nucleoli were stained with anti-nucleolin antibody (NCL, red). Filled arrows indicate strong biotinylation of nucleolar proteins, open arrows highlight nucleoli that are not preferentially labelled in control BirA*-myc expressing cells. (D) Schematic diagram of BirA* fusion constructs used in panels B and C.

